# EEG microstate dynamics during psilocybin intoxication relate to acute experience and persisting psychological changes

**DOI:** 10.64898/2026.06.09.731183

**Authors:** Nikola Jajcay, Čestmír Vejmola, Jakub Korčák, Filip Tylš, Michaela Viktorinová, Vojtěch Viktorin, Anna Bravermanová, Renata Androvičová, Marie Balíková, Jiří Horáček, Martin Brunovský, Jaroslav Hlinka, Tomáš Páleníček

## Abstract

Psilocybin and other serotonergic psychedelics show therapeutic promise for psychiatric disorders, yet objective neural correlates linking the acute psychedelic state to persisting psychological outcomes remain limited. Electroencephalography (EEG) microstate analysis characterizes the rapid spatiotemporal organization of large-scale brain activity, offering a millisecond-resolution window into neural dynamics. Here, we examined resting-state EEG microstates in 15 healthy volunteers who participated in a double-blind, randomized, placebo-controlled crossover study of psilocybin, using both data-driven (three-microstate) and canonical (four-microstate) analysis solutions. EEG was recorded at five time points spanning pre-drug baseline, peak intoxication, and recovery. Psilocybin significantly increased the number of global field power (GFP) peaks and reduced microstate lifespan while increasing frequency of occurrence during peak intoxication (50–100 min post-administration), consistent with accelerated transitions between brain states. Notably, microstate coverage was largely preserved, with only a transient difference at peak intoxication in the 2–20 Hz band-width, suggesting that access to the repertoire of canonical brain states is broadly maintained despite altered temporal dynamics. Critically, individual differences in microstate dynamics during peak intoxication correlated with both acute subjective experience intensity and self-reported psychological changes measured 28 days post-administration, providing exploratory evidence for a link between acute neural dynamics and longer-term experiential outcomes in healthy volunteers. These findings suggest that psilocybin is associated with altered temporal organization of large-scale brain dynamics with largely preserved microstate coverage, and identify EEG microstates as candidate neural markers for psychedelic-induced alterations in consciousness with potential relevance to therapeutic research.

## 1. Introduction

Serotonergic psychedelics, including psilocybin, lysergic acid diethylamide (LSD), and N,N-dimethyltryptamine (DMT), have re-emerged as promising therapeutic tools for psychiatric disorders [1]. After decades of limited research, contemporary clinical trials have demonstrated that psilocybin can induce rapid and sustained therapeutic effects in treatment-resistant depression, anxiety associated with life-threatening illness, and related conditions, often after only one or two administrations [2, 3, 4]. These effects contrast sharply with conventional antidepressants, for which clinically significant effects typically require 6–8 weeks of daily dosing at therapeutically active doses, with only approximately one-third of patients achieving remission on first-line treatment [5] and a substantial proportion experiencing adverse effects that may limit quality of life or result in discontinuation [6]. Instead, psilocybin is associated with a brief but profound alteration of consciousness, accompanied by psychological (i.e., improved mood, well-being, and reduced anxiety) and neural changes that can persist for 3–6 months or longer [7, 8]. Understanding the neural mechanisms that link the acute psychedelic state to enduring clinical improvement has therefore become a central goal of psychedelic neuroscience.

A growing body of evidence indicates that the subjective quality of the acute psychedelic experience plays a critical role in determining therapeutic outcome. Dimensions such as ego dissolution, mystical-type experience, emotional breakthrough, and peak experience consistently predict long-term improvements in mood, well-being, and personality traits in both healthy volunteers and clinical populations [9, 10, 11]. However, reliance on subjective reports alone limits mechanistic inference and clinical translation. Identifying objective neural markers that track the intensity of the psychedelic state and predict long-term outcomes is therefore a priority. Such biomarkers could facilitate patient stratification, optimize dosing strategies, inform mechanism-based models of action, and potentially guide the development of non-pharmacological interventions that emulate therapeutically relevant brain states.

The search for neural predictors of psychedelic efficacy presents unique challenges. Unlike conventional antidepressants, psychedelics exert their primary effects within hours, leaving little opportunity to identify early-response biomarkers based on gradual symptom change. Consequently, research has focused either on baseline predictors—whose identification is complicated by confounds such as medication history and contextual “set and setting”—or on acute neural signatures that may forecast persisting psychological effects. While large-scale datasets required to robustly identify baseline predictors are not yet available, accumulating evidence suggests that acute brain dynamics during the psychedelic state may provide informative markers of therapeutic relevance.

Neuroimaging studies have revealed that psychedelics induce a profound reorganization of large-scale brain networks. Functional magnetic resonance imaging (fMRI) research has shown that psilocybin disrupts the integrity of canonical resting-state networks, particularly the default mode network (DMN), which is implicated in self-referential processing and rumination and shows altered activation patterns in depression [12, 13]. This within-network disintegration is accompanied by increased between-network functional connectivity and reduced hierarchical segregation, resulting in a transiently “hyperconnected” brain state [12]. Converging evidence indicates that psychedelics also increase neural entropy, reflecting a broader and less constrained repertoire of brain states [14, 13, 15]. These changes have been interpreted as reflecting increased neural flexibility and a temporary relaxation of rigid network dynamics that may support psychological change.

Electroencephalography (EEG) studies provide complementary insights at finer temporal resolution. Psilocybin, LSD, and DMT consistently reduce alpha-band power while modulating higher-frequency activity, interpreted as cortical desynchronization and consistent with altered thalamocortical communication [16, 17, 18, 19, 20]. Measures of signal diversity and complexity, including entropy and Lempel-Ziv complexity, reliably increase during the psychedelic state and correlate with subjective experience intensity [21]. Together, these findings suggest that psychedelics accelerate and diversify brain dynamics across multiple temporal scales. However, most EEG studies have focused on spectral or complexity-based metrics, leaving the organization of large-scale spatial dynamics relatively underexplored.

EEG microstate analysis offers a principled framework for characterizing the rapid spatiotemporal organization of brain activity. Microstates are brief periods—typically lasting 60–120 ms—during which the scalp electric field configuration remains quasi-stable before transitioning to a new topography [22, 23]. Often described as the “atoms of thought,” microstates reflect coordinated activity across distributed neural networks. In resting-state EEG, four canonical microstate classes (A–D) typically account for the majority of variance and have been linked to distinct functional networks using simultaneous EEG-fMRI [24]. Microstate analysis yields several interpretable parameters, including duration (lifespan), frequency of occurrence, temporal coverage, and transition structure, which together characterize the temporal syntax of large-scale brain activity.

Alterations in microstate dynamics have been reported across a range of psychiatric and neurological disorders. Schizophrenia and major depression, for example, are associated with reduced microstate duration, altered coverage, and abnormal transition patterns, suggesting that microstates capture clinically meaningful aspects of network dysfunction [25, 26, 27]. More recently, abnormal microstate profiles have been linked to neurodegenerative processes and cognitive decline [28]. These findings position microstates as sensitive markers of large-scale brain dynamics with potential translational relevance. Despite this, the effects of psychedelics on EEG microstate organization have received little systematic investigation.

Given that psilocybin profoundly alters conscious experience, destabilizes canonical network structure, and increases neural entropy, we hypothesized that it would also substantially modulate microstate dynamics. We also explored, in a hypothesis-generating fashion, whether inter-individual differences in microstate dynamics during the acute state would relate to self-reported psychological changes assessed 28 days post-administration.

To test these hypotheses, we examined resting-state EEG microstates in healthy volunteers undergoing placebo and psilocybin sessions. Our primary aims were (i) to characterize the acute effects of psilocybin on microstate temporal organization, and (ii) to determine whether individual differences in microstate dynamics during the psychedelic state relate to both subjective experience and long-term psychological outcomes previously reported in this cohort [11]. By linking millisecond-scale brain dynamics to both phenomenology and persisting effects, this work seeks to evaluate EEG microstates as candidate neural markers of psychedelic-induced changes in consciousness and their potential therapeutic relevance.

## 2. Methods

### 2.1. Study Approval

The study was approved by the local ethics committee of the Prague Psychiatric Centre/National Institute of Mental Health and by the Czech legal authority, the State Institute for Drug Control. It was approved as a clinical trial registered under the EudraCT No. 2012-004579-37. All volunteers signed an informed consent prior to entering the study.

### 2.2. Study Design, Participants, and Recruitment

Data for the present analysis were collected during the first phase of a larger clinical trial consisting of two arms. In each arm, subjects underwent one psilocybin and one placebo session, using a double-blind, randomized, crossover design. While the first arm focused primarily on EEG data collection, the second arm was designed for fMRI data collection. Sessions were separated by at least a 28-day interval. The comprehensive structure of the full trial is detailed in our previous work [11].

Healthy volunteers were recruited via word-of-mouth and community referral. To ensure the exclusion of neurological, psychiatric, or major medical disorders—as well as current or past substance misuse—all participants underwent a rigorous screening process. This included a medical somatoneurological examination, laboratory testing, the Mini-International Neuropsychiatric Interview (M.I.N.I.), and the Minnesota Multiphasic Personality Inventory-2 (MMPI-2). Candidates with a personal or first-degree family history of psychotic illness were excluded. All participants abstained from alcohol and psychoactive substances for at least 48 hours before each session.

For the analyses presented here, we utilized data from the first 20 healthy subjects (8 females; aged 20–35 years). This cohort was recorded using a 21-channel EEG and is identical to the sample described in [29]. Of the 20 enrolled participants, 5 were excluded from the final analyses due to data acquisition problems (N=3 missing sessions; N=2 incomplete EEG recordings due to poor signal quality). All analyses reported here are therefore based on **N = 15** participants.

### 2.3. Drug and Dosage

Participants received a weight-adjusted oral dose of approximately 0.26 mg/kg of pharmaceutical-grade psilocybin encapsulated in gelatin capsules with *Amylum tritici* (wheat starch) as a binder. Placebo capsules contained *Amylum tritici* (wheat starch) only and were visually indistinguishable. Capsules containing either 1 or 5 mg of psilocybin were prepared at the IKEM (Institute of Clinical and Experimental Medicine in Prague) pharmacy. The dose of psilocybin was adjusted by combining capsules containing 1 and 5 mg, increasing or decreasing by 1 mg per 5 kg of body weight, with a 75 kg person receiving 20 mg of psilocybin.

### 2.4. Experimental Design

Each participant completed two experimental days (psilocybin, hereafter *PSI*, and placebo, hereafter *PLA*) in a counterbalanced order. Throughout the dosing sessions, participants were accompanied by a pair of assisting sitters who served supportive roles and actively participated in data collection. A trained nurse/EEG technician was also present throughout the sessions. Upon arrival, baseline physiological measurements and questionnaires were collected. Subjects received an intravenous cannula and were fitted with the EEG gel base cap, and the session began. After receiving the drug or placebo, participants rested comfortably in a reclined position under the continuous supervision of assisting sitters.

Resting-state eyes-closed EEG was recorded at five predefined time points during the whole session: a baseline 10-minute recording acquired before drug ingestion (T1), followed by four 10-minute recordings at 50–60 minutes (T2), 90–100 minutes (T3), 180–190 minutes (T4), and 360–370 minutes (T5) after ingestion. These intervals captured pre-drug baseline, peak and late intoxication, and the post-acute recovery period. No stimulation was presented; participants were instructed to keep their eyes closed and remain still and relaxed. In between the resting-state EEG recordings, subjects listened to music, underwent EEG oddball paradigms [29] and auditory steady-state responses (ASSR) [30], and were repeatedly examined with the Brief Psychiatric Rating Scale (BPRS). Blood samples for psilocin plasma levels were collected at 5 time points (baseline, 1h, 2h, 4h, and 6h after ingestion) throughout the dosing session.

### 2.5. Subjective and Behavioural Measures

Subjective and behavioral effects of psilocybin were assessed using a combination of standardized psychometric instruments that capture acute drug effects, clinical symptomatology during intoxication, and persisting subjective effects after the session.

Acute subjective drug effects were evaluated using the Altered States of Consciousness (ASC) questionnaire [31], which participants completed at the end of each experimental session. The ASC comprises 72 items rated on visual analog scales and yields scores across three main dimensions — Oceanic Boundlessness (OSE), Dread of Ego Dissolution (AIA), and Visionary Restructuralization (VUS), as well as a fourth composite score reflecting the overall intensity of the altered state of waking consciousness (Veränderter Wachbewusstseinszustand, VWB), providing a multidimensional characterization of the psychedelic experience.

Objective clinical and behavioral changes during the session were assessed using the BPRS [32]. The BPRS was administered repeatedly over the course of each session by assisting sitters, specifically before drug/placebo intake (T=0 min) and at 70 and 180 minutes after ingestion. The scale rates 18 psychiatric symptoms, each on a 7-point scale (0–6, ranging from “not present” to “very severe”), which can be clustered into five symptom domains: FI (anxiety, depression), FII (withdrawal, retardation), FIII (thought disturbance and hallucinations), FIV (tension, excitement), and FV (hostile suspiciousness), as described in [29].

Persisting subjective effects were assessed 28 days after each experimental session [11] using the Persisting Effects Questionnaire (PEQ) [10, 33]. The PEQ captures persisting subjective changes attributed to the psychedelic experience, including alterations in mood, attitudes, behavior, well-being, and perceived personal meaning. Items are rated on Likert-type scales and summarized into positive and negative persisting-effect domains, allowing quantification of persisting experiential impact beyond the acute intoxication phase.

### 2.6. Psilocin Plasma Levels

Blood samples were drawn via an indwelling cannula inserted before the start of each session. The samples were collected at 5 time points: before ingestion (baseline) and at 1, 2, 4, and 6 hours after ingestion of the capsules. Blood samples were centrifuged at room temperature for 10 min at 4000 rpm. The separated sera were then stored at *−*20 °C until analyzed. Psilocin in sera was analyzed by gas chromatography/mass spectrometry (GC-MS). The analysis itself was performed on a GC HP model 6890 A with 5973 MSD and capillary HP5-MS. For details on the analyses, see our previous work [29].

### 2.7. EEG Recording

EEG signals were recorded using a 21-channel Ag/AgCl cap arranged according to the extended 10–20 system (original 10–20 nomenclature; ElectroCap International, USA). Four bipolar EOG channels monitored eye movements. Signals were amplified with a BioSDA09 digital EEG amplifier (M&I, Prague, Czech Republic), sampled at 1000 Hz. Electrode impedances were kept below 5 kΩ. All recordings were obtained in an electrically shielded, sound-attenuated room with participants resting in a reclined position.

### 2.8. EEG Data Preprocessing

EEG preprocessing was performed in *BrainVision Analyzer 2*.*1*.*1* (Brain Products GmbH, Gilching, Germany). Raw data were visually and semi-automatically inspected for artifacts.

Signals were filtered using zero-phase infinite impulse response (IIR) Butterworth filters (0.5–100 Hz band-pass, 8th order) together with a 50 Hz notch filter. Independent component analysis (ICA; FastICA, 24 components; 21 scalp EEG channels and four EOG channels used as input) was applied to the continuous data. Artifactual ICA components, primarily those reflecting ocular activity, were identified semi-automatically and removed via inverse ICA before data reconstruction.

Following ICA cleaning, EEG was re-referenced to the average of 19 scalp electrodes (Fp1, Fp2, F3, F4, F7, F8, Fz, C3, C4, Cz, T3, T4, T5, T6, P3, P4, Pz, O1, O2). The data were then downsampled to 250 Hz using spline interpolation after anti-alias filtering at 112.5 Hz (24 dB/oct).

Continuous EEG recordings were segmented into non-overlapping 2-second epochs, with all previously marked artifact intervals excluded. Only clean epochs were retained for microstate analysis. Preprocessed EEG was exported in ASCII format for subsequent computational processing in Python.

### 2.9. Microstate analysis

EEG microstate analysis was performed for each participant and condition separately using Python. Within microstate analysis, the multichannel EEG signal is viewed as a series of instantaneous topographies of potential. We extracted the global field power (GFP) peaks in order to account only for temporal points with the highest signal-to-noise ratio [22, 34]. The GFP at each temporal instant is equal to the root mean square across the average-referenced electrodes (i.e. a standard deviation of the signal):

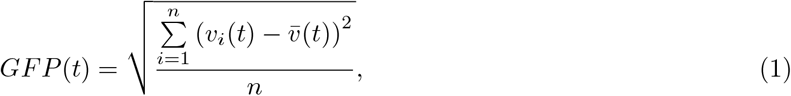

with *v*_*i*_(*t*) being the voltage at electrode *i* at time instant *t*, 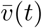 is the mean voltage across all electrodes in a given time instant *t*, and *n* is the number of electrodes. Scalp potential topographies (or maps) at GFP time-series maxima represent the highest field strength and the greatest signal-to-noise ratio. The microstate analysis then includes two steps: first, identifying a set of microstate maps; second, projecting the original multichannel EEG recording onto the basis of microstate maps, thereby converting the EEG signal into a sequence of microstate maps (see Supplementary Figure 10 for an overview).

The maps at the maxima of GFP time-series were submitted into a modified hierarchical clustering algorithm [35]. Briefly, all maps are initially considered as independent clusters and in each iteration, the worst cluster is identified and dissolved; its constituent maps are redistributed to the remaining clusters according to the strongest Pearson product-moment correlation in the form

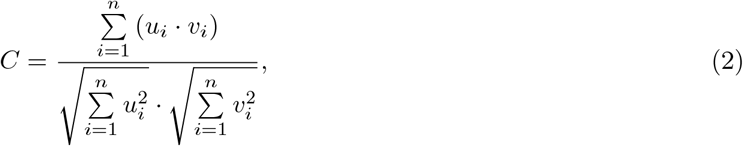

where sums are taken over *i* electrodes and *u* and *v* represent two topographies. These steps are repeated until the desired number of clusters remains (see e.g. [36]).

We used the above-described microstate algorithm to cluster original maps from each participant and condition into microstate maps, specific for each subject and condition. Next, in order to obtain *condition-specific* maps, we submitted maps from each participant from given condition to another round of clustering separately for each condition, thus obtained mean maps for given condition. All maps (both individual and *condition-specific*) were labeled class A, B, C, and D based on resemblance (i.e. Pearson product-moment correlation) to normative template maps following Koenig et al. [37] (Supplementary Figure 10).

After finding canonical microstate maps for each subject and each condition, the original multichannel EEG recording was transformed into a temporal sequence of microstate maps by the following procedure: one of the canonical maps is assigned for each GFP peak based on the highest absolute Pearson product-moment correlation (i.e., map polarity is disregarded) and if two subsequent GFP peaks are assigned different maps, we identified the temporal midpoint and created a microstate transition there. No minimum segment duration threshold or temporal smoothing was applied during back-projection. From the sequence of microstate maps, we calculated the following features:

#### Average lifespan of microstates

Lifespan of microstates is calculated as the time during which all successive original topographies were assigned the same microstate class [25] (cf. Supplementary Figure 10).

#### Microstate coverage

The coverage of a microstate is calculated by taking a ratio of the total time spent in that particular microstate over total recording time [25].

#### Frequency of occurrence

Microstate frequency is determined by counting unique appearances of each microstate in one second of the recording [25].

#### Microstate transition probabilities

The number of transitions from each microstate class into other microstate class is counted and normalised to fractions of all between-class transitions, yielding a *N ×N* matrix with zero diagonal of transition probabilities [25]. Transition probabilities were computed for completeness but are not analyzed statistically in the present study, as the primary focus is on temporal (lifespan, frequency of occurrence) and spatial (coverage) microstate parameters.

After the EEG data were preprocessed, divided into epochs, and artifacts removed, we were left with 20 epochs of 2 s duration, totaling 40 seconds of the signal per participant, per condition (placebo: *PLA* vs. psilocybin: *PSI*), and time (*T1* through *T5*). We then submitted these data to the microstate analysis described above. All our subsequent analyses rely on two different filtering approaches: since participants in our study were administered psilocybin, which shifts relative spectral maximum towards faster frequencies, we opted for a wider bandwidth, that is 1–40 Hz. On the other hand, in order to be able to compare our results with the existing body of literature, we repeated all analyses with a more “traditional” microstate filtering option, that is 2–20 Hz [37, 38, 39].

A fundamental methodological decision concerned the optimal number of microstates to extract. The cross-validation (CV) criterion [40] indicated three microstates as optimal; however, the canonical approach in the microstate literature employs four microstates [23]. As noted by Murray et al. [40], the CV criterion can underestimate the true number of classes in datasets with lower electrode density, and a recent rat study similarly found it to favour three microstates [41]. We therefore analysed both three- and four-microstate solutions across both frequency bands, reporting the 1–40 Hz / three-microstate solution as primary (supported by the CV criterion) and the 2–20 Hz / four-microstate solution as secondary (following traditional methodology).

### 2.10. Statistical analyses

To compare the number of GFP peaks across times and conditions, we used repeated-measures ANOVA, followed by pairwise repeated-measures t-test post hoc with the correction for multiple comparison using the Benjamini-Hochberg step-up procedure to control the False Discovery Rate (FDR). Similar procedure (repeated-measures ANOVA + pairwise repeated-measures t-test post hoc) was employed to compare microstate features (lifespan, coverage, and frequency of occurrence).

To assess relationships between subject experience data, we used Spearman’s rank correlation coefficient *ρ* to mitigate non-normality, with significance assessed using the Benjamini-Hochberg FDR procedure to control for multiple comparisons across all correlation pairs within each matrix. To minimize the number of correlations between experience data and microstate features and streamline the comparison, we aggregated both the experience data and microstate features using principal component analysis (PCA) and used the first PCA component, explaining the largest proportion of variance in the whole dataset. The first PCA component in the case of microstate features explains approximately 80% of the variance, whereas in the case of experience data it ranges between 54 and 72%.

## 3. Results

### 3.1. Microstate analysis

#### 3.1.1. Global field power and ideal number of microstates

First, we compared the number of peaks in GFP curves between conditions. Our hypothesis was that the number of peaks in the psilocybin condition would be higher for relevant times (i.e. *T2, T3*, and, possibly, *T4*), since classical psychedelics tend to shift spectral maxima towards higher frequencies, hence the signal itself is faster and therefore we expected the number of peaks to be higher. This was validated using repeated measures ANOVA which identified **condition** as a significant factor in 2–20 Hz bandwidth, while all (**condition, time**, and their **interaction**) factors were regarded as significant in 1–40 Hz bandwidth. Figure 1 illustrates this point: in the right-hand side of the figure (1–40 Hz bandwidth) the number of GFP peaks in psilocybin condition is indeed significantly higher than placebo in *T2, T3*, and *T4* times. The number of GFP peaks for psilocybin *T3* condition is significantly higher in 2–20 Hz bandwidth as well (left-hand side of Figure 1). Figure 1 also indicates significant differences as revealed by pairwise t-tests with Benjamini-Hochberg procedure [42] employed in order to control false discovery rate (FDR). Higher number of GFP peaks results in higher number of topographic maps submitted into the microstate clustering algorithm.

**Figure 1:**
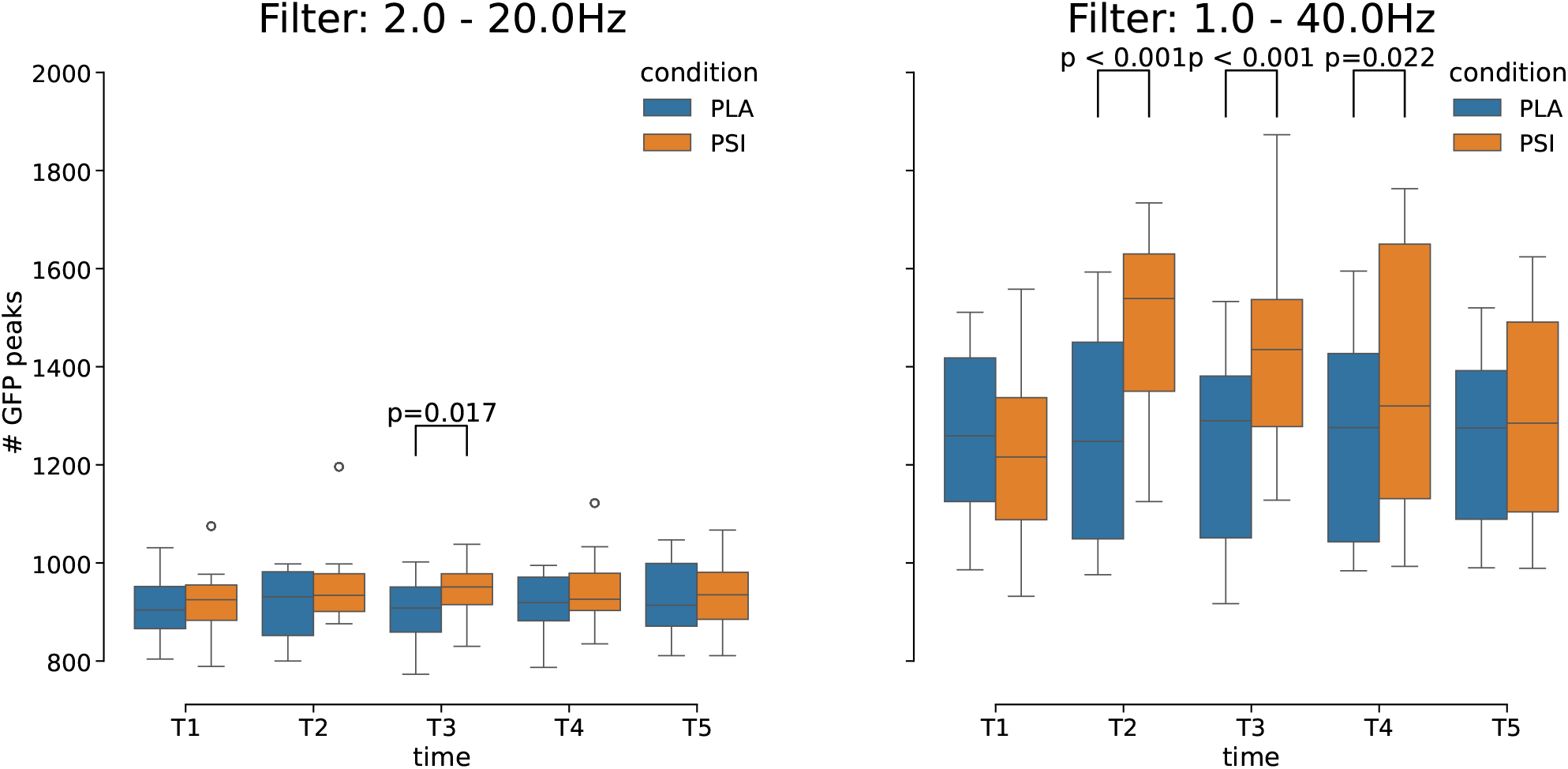
Box plots showing number of GFP peaks in 2 different conditions (placebo and psilocybin) and 5 different acquisition times. The number of GFP peaks is higher for psilocybin in *T2, T3*, and *T4* in both bandwidths, albeit *significantly* higher only in *T3* 2–20 Hz bandwidth and in *T2, T3*, and *T4* in 1–40 Hz bandwidth. *p-values* are based on *pairwise t-test* post hoc (preceded by repeated measures ANOVA) and are corrected for multiple comparisons using *Benjamini-Hochberg* FDR procedure.

Traditionally, the microstate studies compute 4 canonical microstates and refer to the seminal early works in this area (e.g. Lehmann et al. [22] or Pascual-Marqui et al. [43]). Given that microstate analysis has received limited systematic investigation in the context of psychedelic EEG [19], we also tested the ideal number of microstates as obtained from the cross-validation (CV) test proposed by Pascual-Marqui et al. [43]. Cross-validation criterion states that the ideal number of microstates is the number *q* which minimises modified cross-validation variance 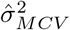 given by the following function [43]:

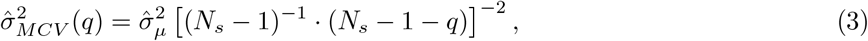

where 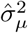 is the microstate model nonpredictive residual variance for *q* different microstates and *N*_*s*_ is the number of electrodes.

We computed the 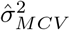 curve for each data snippet with changing number of states *q* ∈ {1, …, 10} in all times and conditions and then found the minimum within each curve. Figure 2 summarises the ideal number of states (*q* that minimises the 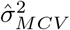 curve) in our dataset. Unexpectedly, the ideal number of microstates in each condition and time across the subject (aggregating with median) was 3. Overall, we observe the trend towards lower number of states in wider, 1–40 Hz, bandwidth, however the ideal number of states was 3 for both bandwidths.

**Figure 2:**
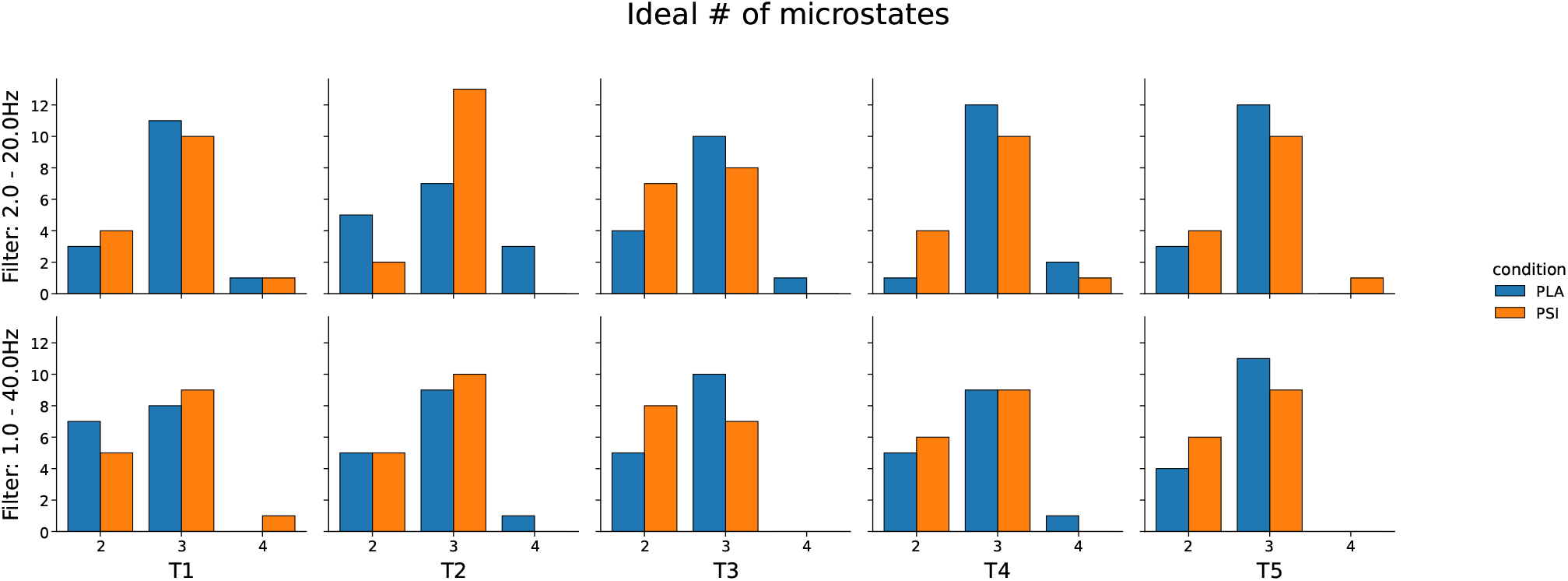
Ideal number of microstates based on variance test according to Pascual-Marqui et al. [43]. First row shows 2–20 Hz bandwidth, while the second row shows wider, 1–40 Hz, bandwidth. The individual columns stand for times *T1* through *T5*, while the colour encodes condition (*PLA* vs. *PSI*). The results indicate that ideal number of microstates (that is the number which minimises modified cross-validation variance 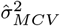) is **3** in both bandwidths, both conditions, and all times.

Based on these results, we decided to continue our analyses within two branches: the main branch is 1– 40 Hz bandwidth with decomposition into 3 canonical microstates, while the second is “classical” microstate analysis, that is 2–20 Hz bandwidth with 4 canonical states.

#### 3.1.2. Microstate maps

After initial EEG signal analysis with respect to the number of GFP peaks and ideal number of microstates within each bandwidth, we present the actual microstate topographies in Figures 9 and 8 in supplementary information. The *time-condition* specific topographies, i.e. the average microstates per condition and time, were obtained firstly by running microstate clustering algorithm on all our data (that is for each participant, each condition, and each time) and all maps from one group were subsequently submitted to another round of clustering.

The maps for 1–40 Hz bandwidth, for which the ideal number of microstates was 3 (cf. Figure 2), are shown in Figure 8 in supplementary information for each of the *time-condition* combination. Below each map we also report the Pearson product-moment correlation (see eq. 2) between particular map and microstate “template maps” provided by Koenig et al. [37]. These template maps serve as the gold standard in microstate analysis. The correlations indicate (group-level mean Pearson product-moment correlations per bandwidth, per condition, and per time are all *C >* 0.8; individual maps may fall below this threshold) that all our maps resemble template maps to a high degree and therefore we are able to take the analysis to the next step and compute some microstate statistics using these canonical maps.

Similarly as for wider bandwidth, the 2–20 Hz bandwidth maps (see Figure 9 in supplementary information) were compared with template maps from Koenig et al. [37]. We conclude that in 2–20 Hz bandwidth all our maps bear a resemblance to template and the overall correlation is even higher than in the 1–40 Hz case. Consequently, we take also this branch of our analysis to the next step and submit the data to microstate statistics computation.

We also tested whether there is a significant difference in explained variance in GFP peaks by our microstate decomposition. In the 2–20 Hz bandwidth with 4 canonical states there was no significant difference between times (*F* (4, 56) = 2.042, *p* = 0.127), condition (*F* (1, 14) = 4.226, *p* = 0.059), nor their interaction (*F* (4, 56) = 1.963, *p* = 0.146) as indicated by 2-way repeated measures ANOVA (p-values are Greenhouse-Geisser-corrected for sphericity). However, in the 1–40 Hz bandwidth with 3 canonical microstates, repeated measures ANOVA showed significant dependence on condition with *F* (1, 14) = 5.675, *p* = 0.032, where the variance explained in placebo condition was slightly higher (0.707 *±*0.065, mean *±* SD) than in the psilocybin condition (0.682 *±* 0.055) (unbiased Cohen’s *d* = 0.414). Other factors were not identified as significant (time: *F* (4, 56) = 2.521, *p* = 0.074 and interaction condition*×* time: *F* (4, 56) = 2.751, *p* = 0.066).

#### 3.1.3. Microstate features

After performing base analysis of GFP peaks, cross-validation test for ideal number of canonical maps and visualising the canonical maps themselves for both our filtering paradigms we computed typical microstate features which are the average lifespan of microstates, microstate coverage, frequency of occurrence, and their transition probabilities.

The results of average microstate lifespan for both filtering paradigms are shown in Figure 3 while the significant differences in average lifespan per conditions, microstate, and time are described in Table 1. Generally, and in line with our initial hypothesis, the significant differences between placebo and psilocybin conditions are mainly in the *T2* and *T3* times, when the effect of the intoxication is the strongest with psilocybin condition tending to lower average lifespan, consistent with faster state switching. This is true for both filtering paradigms. Similar results (faster state switching) were observed in interaction with particular microstates: microstates B and C in 1–40 Hz bandwidth. The effect sizes as measured with Cohen’s *d* were strong in this interaction setting (cf. Table 1; Supplementary Figure 11). Lastly, even individual microstate factor while disregarding time factor was significantly different in the 1–40 Hz filtering paradigm (microstates A, B, and C).

**Table 1:** Significant differences in lifespan between placebo and psilocybin condition. We show difference in various combinations with two factors (microstate, time) and their interaction. All statistics are based on *pairwise t-test* post hoc (preceded by repeated-measures ANOVA), *p-values* are corrected for multiple comparisons using *Benjamini-Hochberg* FDR procedure, and **d** denotes unbiased Cohen’s *d* effect size.

| Filter: 2–20 Hz |  |  |  |  | Filter: 1–40 Hz |  |  |  |  |
| --- | --- | --- | --- | --- | --- | --- | --- | --- | --- |
| microstate | time | T | p-value | d | microstate | time | T | p-value | d |
| – | T2 | 2.989 | 0.037 | 0.975 | A | – | 2.147 | 0.050 | 0.481 |
| – | T3 | 2.774 | 0.037 | 0.730 | B | – | 3.370 | 0.007 | 0.481 |
|  |  |  |  |  | C | – | 3.524 | 0.007 | 0.587 |
|  |  |  |  |  | – | T2 | 3.369 | 0.006 | 1.196 |
|  |  |  |  |  | – | T3 | 3.897 | 0.006 | 0.978 |
|  |  |  |  |  | B | T2 | 3.390 | 0.018 | 0.958 |
|  |  |  |  |  | B | T3 | 2.729 | 0.041 | 0.710 |
|  |  |  |  |  | C | T2 | 3.642 | 0.013 | 1.180 |
|  |  |  |  |  | C | T3 | 3.124 | 0.019 | 0.868 |

**Figure 3:**
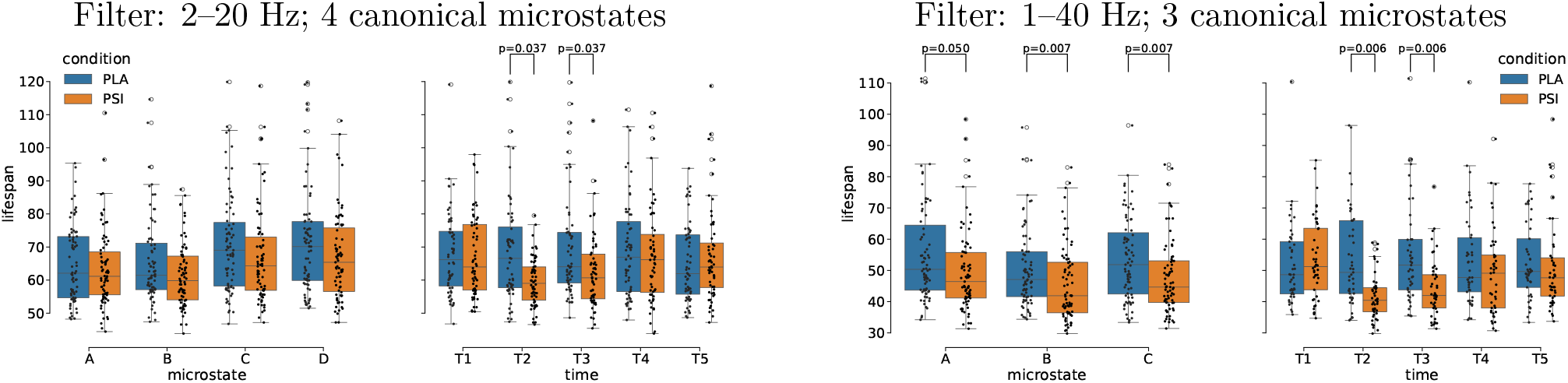
Box plots of average microstate lifespan in ms. For each filtering paradigm (left: 2–20 Hz; right: 1–40 Hz), lifespans are grouped by microstate (left sub-panel) and time (right sub-panel); interaction effects are shown in Supplementary Figure 11. Significant differences between placebo and psilocybin condition are marked with vertical lines and their respective *p-values* as based on *pairwise t-test* post hoc (preceded by repeated measures ANOVA) and are corrected for multiple comparisons using *Benjamini-Hochberg* FDR procedure. For tabulated results please refer to Table 1.

Similarly to the lifespan, coverage results are depicted in Supplementary Figure 12. In the 2–20 Hz bandwidth, a significant difference in coverage between conditions was found at *T2* (pairwise t-test post hoc, *p* = 0.015, Benjamini-Hochberg FDR corrected). No significant differences were found for any other time or microstate combination, nor in the 1–40 Hz bandwidth. Overall, as the figure shows, the means of coverage were similar across both factors and their interaction, while the interquartile ranges were narrower in the psilocybin condition, particularly at *T2* and *T3*.

Our final measure of interest was the frequency of occurrence. The main results are plotted in Figure 4. Since frequency of occurrence measures similar underlying dynamics as the average lifespan (the relationship between the two is inverse), we expected similar outcomes as with lifespan. As expected, the main differences between two conditions are observed in *T2* and *T3* times; moreover, the effect still persists into *T4* in the 1–40 Hz filtering paradigm. Similar results were observed in interaction with particular microstates: all three microstates in the 1–40 Hz bandwidth and microstate A in 2–20 Hz bandwidth. The effect sizes as measured with Cohen’s *d* were strong in this interaction setting (cf. Table 2). Lastly, even individual microstate factor while disregarding time factor was significantly different in the 1–40 Hz filtering paradigm (all three microstates A, B, and C).

**Table 2:** Significant differences in frequency of occurrence between placebo and psilocybin condition. We show difference in various combinations with two factors (microstate, time) and their interaction. All statistics are based on *pairwise t-test* post hoc (preceded by repeated-measures ANOVA), *p-values* are corrected for multiple comparisons using *Benjamini-Hochberg* FDR procedure, and **d** denotes unbiased Cohen’s *d* effect size.

| microstate | time | T | p-value | d |
| --- | --- | --- | --- | --- |
| – | T2 | 3.184 | 0.022 | 1.000 |
| – | T3 | 3.045 | 0.022 | 0.709 |
| A | T2 | 3.623 | 0.014 | 1.158 |

Table 2: Significant differences in frequency of occurrence between placebo and psilocybin condition.
| microstate | time | T | p-value | d |
| --- | --- | --- | --- | --- |
| A | – | 5.535 | < 0.001 | 0.613 |
| B | – | 3.712 | 0.002 | 0.526 |
| C | – | 3.876 | 0.002 | 0.640 |
| – | T2 | 4.354 | 0.002 | 1.284 |
| – | T3 | 5.319 | < 0.001 | 1.028 |
| – | T4 | 2.480 | 0.044 | 0.409 |
| A | T2 | 5.384 | < 0.001 | 1.445 |
| A | T4 | 2.881 | 0.030 | 0.568 |
| B | T2 | 2.941 | 0.027 | 1.088 |
| B | T3 | 5.583 | < 0.001 | 1.047 |
| C | T2 | 3.180 | 0.017 | 0.966 |
| C | T3 | 4.259 | 0.004 | 1.101 |

**Figure 4:**
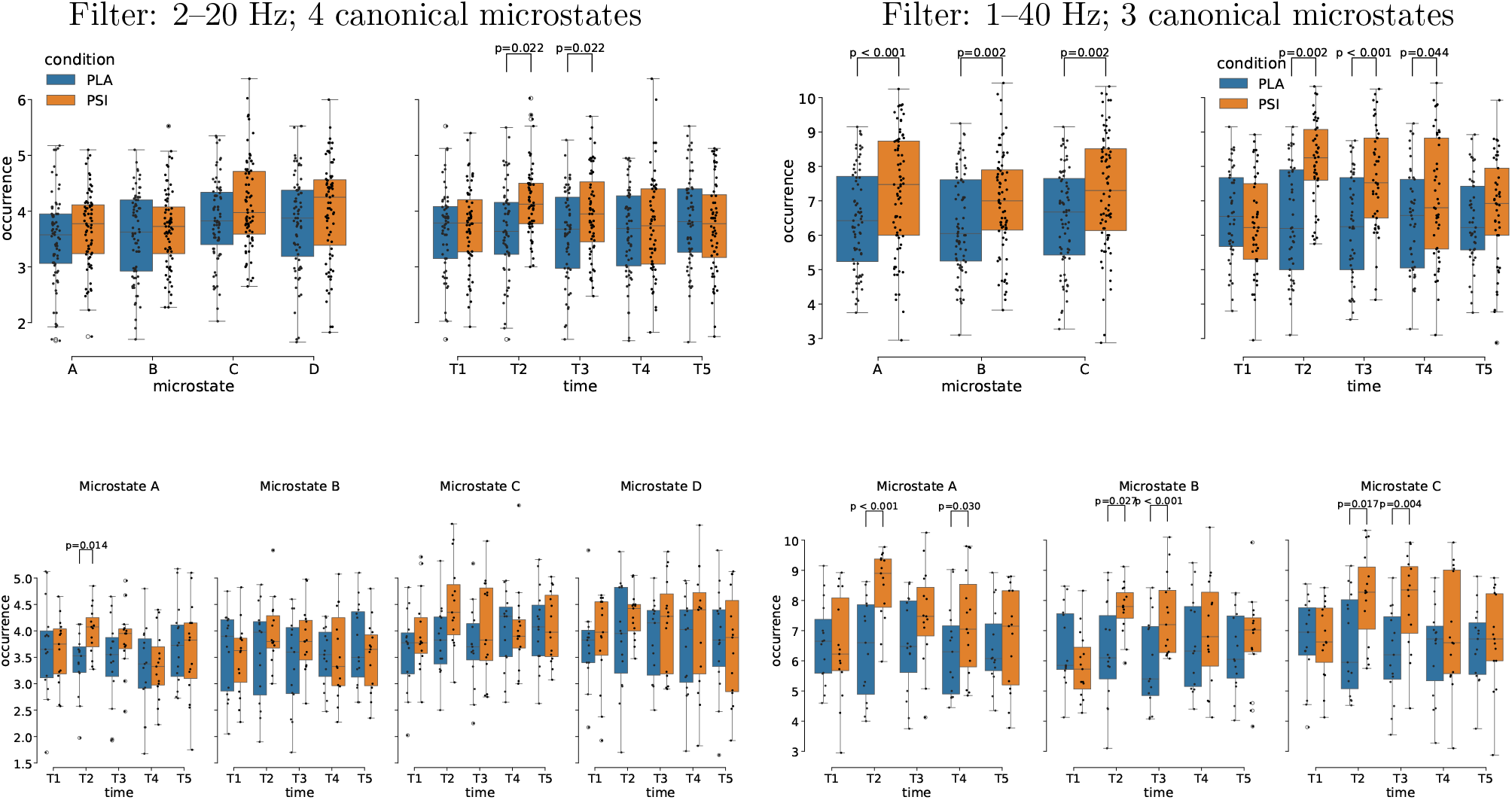
Box plots of frequency of occurrence for microstates per second. For each filtering paradigm (left: 2–20 Hz; right: 1–40 Hz), frequency of occurrence values are grouped by microstate (left sub-panel) and time (right sub-panel), with their interaction shown in the bottom row. Significant differences between placebo and psilocybin condition are marked with vertical lines and their respective *p-values* as based on *pairwise t-test* post hoc (preceded by repeated-measures ANOVA) and are corrected for multiple comparisons using *Benjamini-Hochberg* FDR procedure. For tabulated results please refer to Table 2.

### 3.2. Relationship between microstates and psychedelic phenomenology

Given the pronounced effects of psilocybin on microstate temporal dynamics, we next examined whether these neural changes relate to subjective experience and clinical ratings collected during the experimental sessions. As expected given the established psychotropic effects of psilocybin, all four ASC dimensions were significantly elevated in the psilocybin condition compared to placebo: Oceanic Boundlessness (OSE: *t*(14) = 15.516, *p* =*<* 0.001, *d* = 5.045), Dread of Ego Dissolution (AIA: *t*(14) = 7.949, *p* =*<* 0.001, *d* = 2.842), Visionary Restructuralization (VUS: *t*(14) = 19.254, *p* =*<* 0.001, *d* = 7.072), and the overall altered state of waking consciousness composite score (VWB: *t*(14) = 15.646, *p* =*<* 0.001, *d* = 5.538).

Similarly, the Brief Psychiatric Rating Scale revealed significant condition-related differences across most symptom domains during peak intoxication. At baseline (*T=0min*), no differences were observed between conditions. However, at 70 and 180 minutes post-administration, BPRS factors showed substantial elevations under psilocybin, particularly in thought disturbance and hallucinations (FIII), withdrawal and retardation (FII), and tension and excitement (FIV); anxiety and depression (FI) showed no significant difference at either time point; hostile suspiciousness (FV) was significantly elevated at T=70 min (*p* = 0.033, *d* = 0.861) but not at T=180 min (*p* = 0.164). Full statistical details are reported in Table 3.

**Table 3:** Mean differences in five factors of the Brief Psychiatric Rating Scale (BPRS) at two post-ingestion time points (T=0 min baseline not shown as no differences were observed) between two experimental conditions: placebo and psilocybin. Shown are *t statistic* values and their respective *p-values* and Cohen’s *d* effect sizes as computed from the paired t-test (*df* = 14). Scale labels BP-5:FKT-I through BP-5:FKT-V correspond to factors FI through FV as defined in the main text.

| scale | T | p-value | d |
| --- | --- | --- | --- |
| BP-5:FKT-I | 1.103 | 0.288 | 0.335 |
| BP-5:FKT-II | 4.680 | < 0.001 | 1.507 |
| BP-5:FKT-III | 6.964 | < 0.001 | 2.543 |
| BP-5:FKT-IV | 3.415 | 0.004 | 1.255 |
| BP-5:FKT-V | 2.358 | 0.033 | 0.861 |

Table 3: Mean differences in five factors of the Brief Psychiatric Rating Scale (BPRS) at two post-ingestion time points (*T*=0min baseline not shown as no differences were observed) between two experimental conditions: placebo and psilocybin.
| scale | T | p-value | d |
| --- | --- | --- | --- |
| BP-5:FKT-I | 1.625 | 0.126 | 0.587 |
| BP-5:FKT-II | 3.560 | 0.003 | 1.304 |
| BP-5:FKT-III | 6.086 | < 0.001 | 2.222 |
| BP-5:FKT-IV | 3.224 | 0.006 | 1.177 |
| BP-5:FKT-V | 1.468 | 0.164 | 0.536 |

To examine how self-reported subjective experiences (as captured by the ASC scales) relate to observer-rated clinical symptoms (as captured by the BPRS) during the psilocybin session, we computed a correlation matrix including psilocin concentration in the blood at 5 different times during the experiment and the dose in mg per participant. The matrix is shown in Figure 5. As clearly seen from the correlation matrix, individual types of measures (ASC, BPRS at three different times) are all internally consistent, hence they create clusters of high positive correlation in the grand matrix. Moreover, ASC scales correlate with BPRS scales at 70 and 180 minutes into the experiment, and all scales correlate with psilocin concentration in the blood, in particular after 60 and 120 minutes from the start of the experiment. This confirmed that participants experienced a robust altered state of consciousness, as reflected across both self-reported and observer-rated measures and in plasma psilocin concentration.

**Figure 5:**
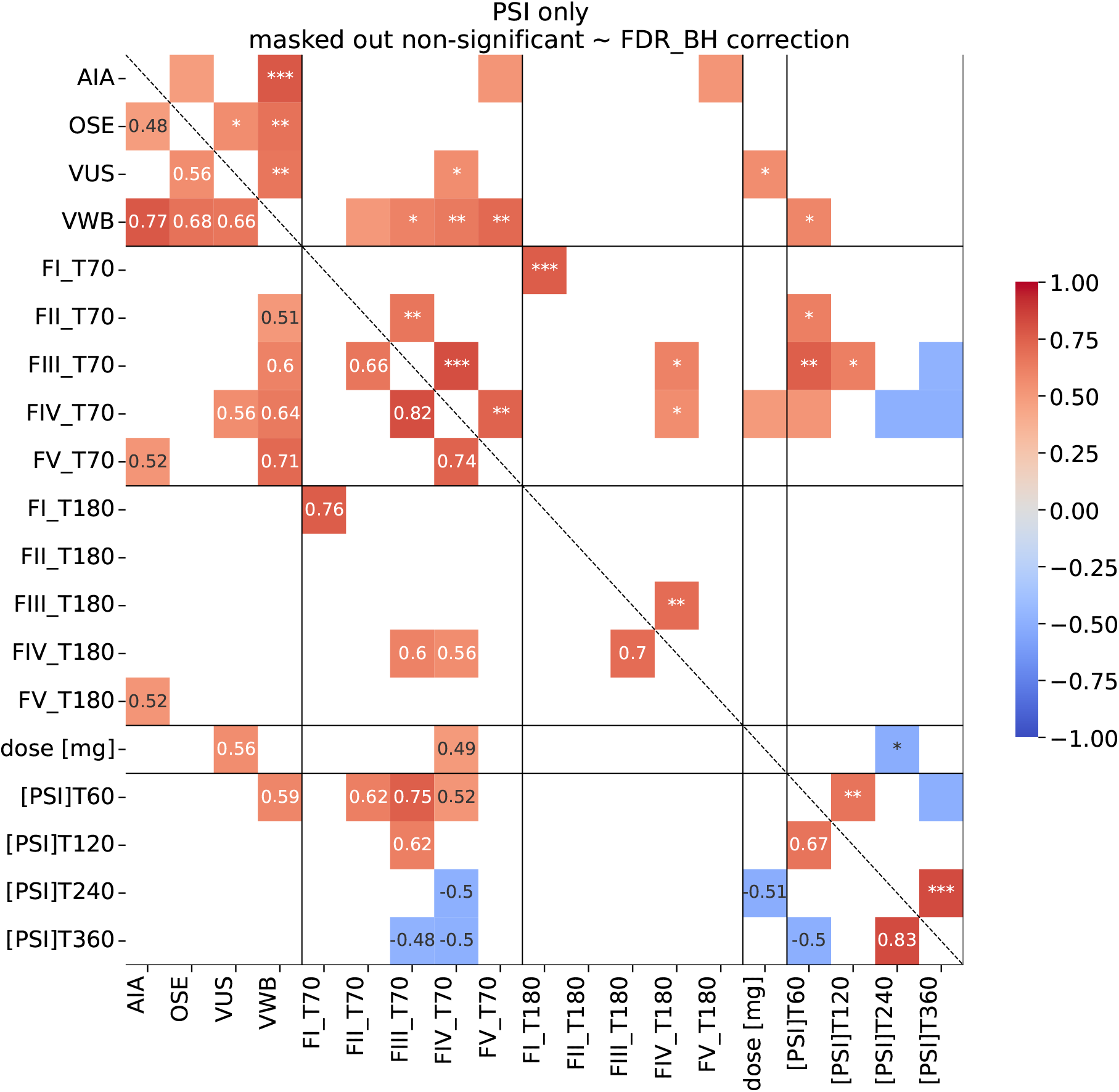
Correlation matrix of measured experience data during the psilocybin experiment. First four measures (AIA = Dread of Ego Dissolution; OSE = Oceanic Boundlessness; VUS = Visionary Restructuralization; VWB = Veränderter Wachbewusst-seinszustand, i.e., altered state of waking consciousness) decode altered states of consciousness (*ASC*) subscales. BPRS factors (FI–FV) are derived from the Brief Psychiatric Rating Scale [29]: FI = anxiety/depression, FII = withdrawal/retardation, FIII = thought disturbance/hallucinations, FIV = tension/excitement, FV = hostile suspiciousness; subscript denotes assessment time (70 or 180 min). [PSI]*T*_*xx*_ denotes plasma psilocin concentration at time *xx* minutes post-administration; *dose* shows psilocybin dosage in mg. Only statistically significant correlations (Benjamini-Hochberg FDR corrected) are shown: the lower triangle displays Spearman *ρ* values; the upper triangle displays significance markers (*\*p <* 0.05, *\*\* p <* 0.01, *\* * * p <* 0.001). Non-significant cells are left blank.

To identify which (if any) microstate feature relates to the changes in psychedelic scales, we plotted Figure 6 as the correlation matrix between aggregated scales (means of ASC, BPRS at relevant times — 70 and 180 minutes of the experiment), dose, and psilocin concentration on the one hand and aggregated microstate features on the other. Aggregation via PCA followed the same procedure described in the Statistical Analyses section.

**Figure 6:**
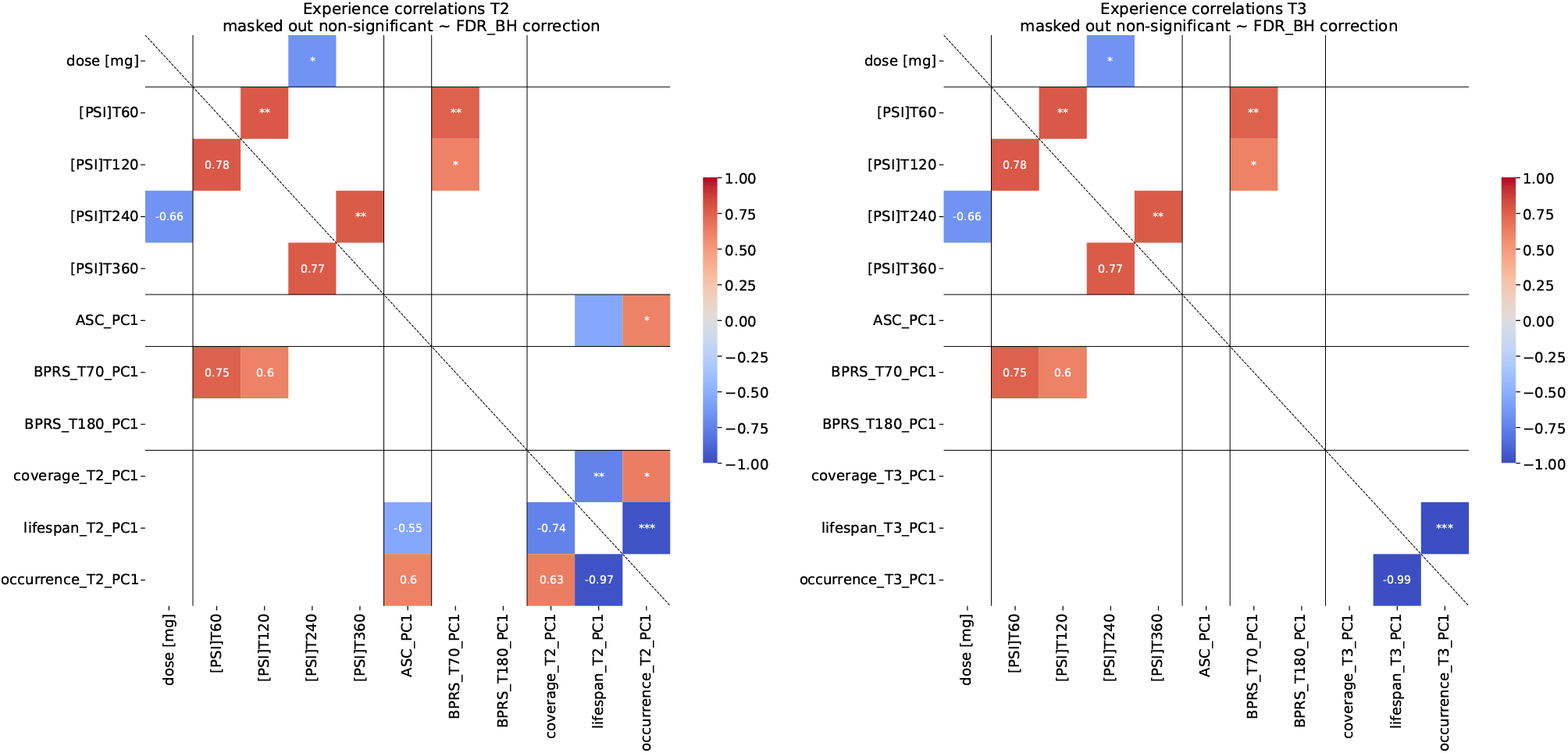
Correlation matrix of experience data (ASC and BPRS questionnaires, psilocin concentration, and psilocybin dose) and aggregated microstate features. Left-hand side shows experience correlations with microstate features at the T2 time, while the right-hand side shows correlations at the T3 time. More details on microstate features aggregation in the main text. Only statistically significant correlations (Benjamini-Hochberg FDR corrected) are shown: the lower triangle displays Spearman *ρ* values; the upper triangle displays significance markers (*\* p <* 0.05, *\*\*p <* 0.01, *\* * *p <* 0.001). Non-significant cells are left blank.

To assess whether acute microstate changes relate to persisting psychological effects, we examined correlations between microstate features during peak intoxication (T2, T3) and PEQ scores obtained 28 days post-administration. As shown in Figure 7, the first PCA component of microstate features correlated with positive persisting effects, particularly for the T2 time point. Notably, this relationship was specific to positive persisting effects; negative persisting effects showed low overall scores (mean negative PEQ score *<* 10% of scale maximum) and no significant associations with microstate dynamics.

**Figure 7:**
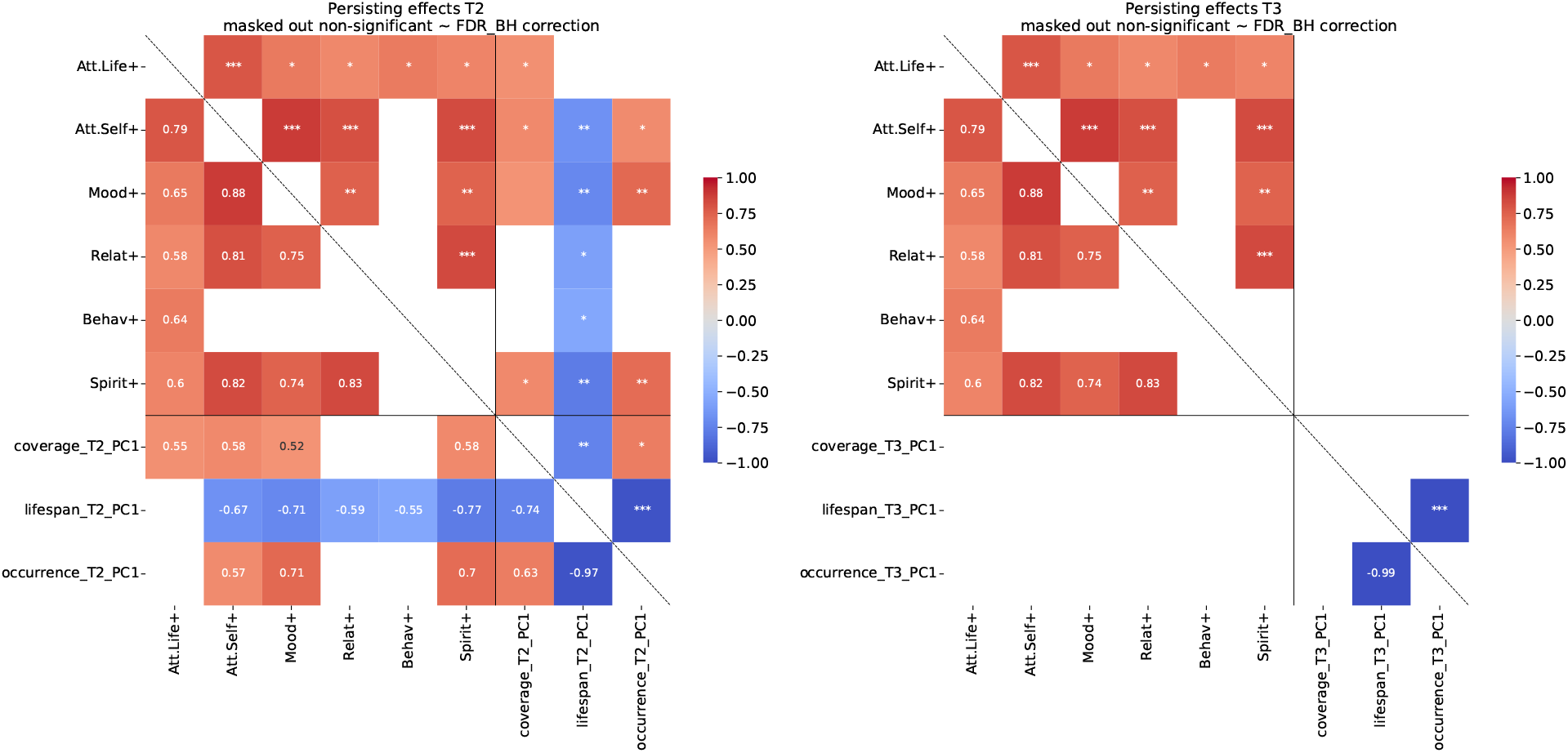
Correlation matrix of persisting effects questionnaire and aggregated microstate features. Left-hand side shows persistent effect correlations with microstate features at the T2 time, while the right-hand side shows correlations at the T3 time. Abbreviated PEQ domain labels: Att.Life+ = Attitudes about Life (positive change); Att.Self+ = Attitudes about Self (positive change); Mood+ = Mood Changes (positive change); Relat+ = Relationships (positive change); Behav+ = Behavioral Changes (positive change); Spirit+ = Spirituality (positive change). More details on microstate features aggregation in the main text. Only statistically significant correlations (Benjamini-Hochberg FDR corrected) are shown: the lower triangle displays Spearman *ρ* values; the upper triangle displays significance markers (*\* p <* 0.05, *\*\* p <* 0.01, *\* * *p <* 0.001). Non-significant cells are left blank.

**Figure 8:**
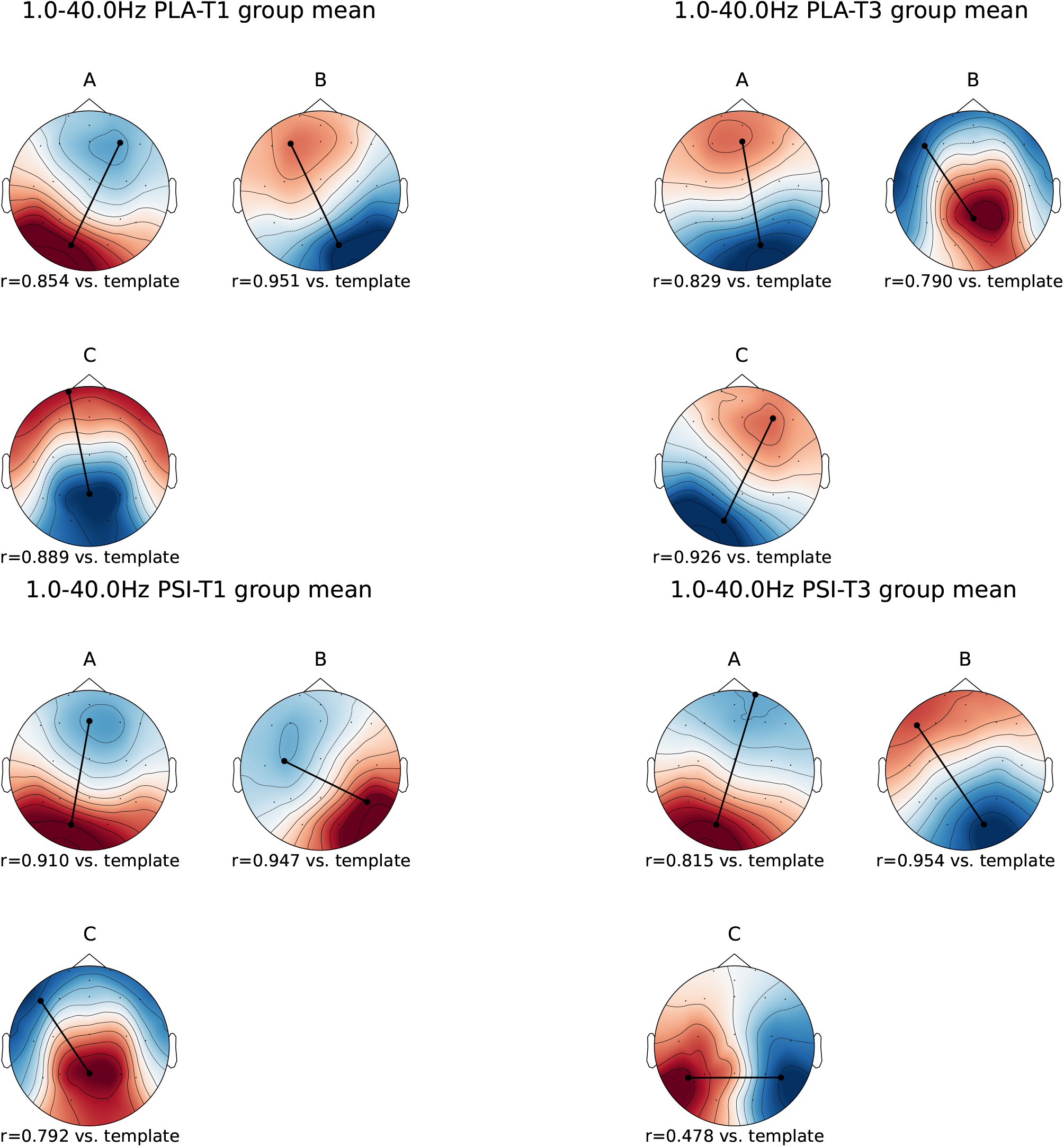
Condition specific topographies of microstates **A, B**, and **C** as found by the microstate algorithm for 1–40 Hz bandwidth. Top row: mean maps for placebo T1 (left) and placebo T3 (right); bottom row: mean maps for psilocybin T1 (left) and psilocybin T3 (right). The correlations (*r*) below the plots signify spatial correlation (cf. eq. 2) between each map and template maps provided by Koenig et al. [37].

**Figure 9:**
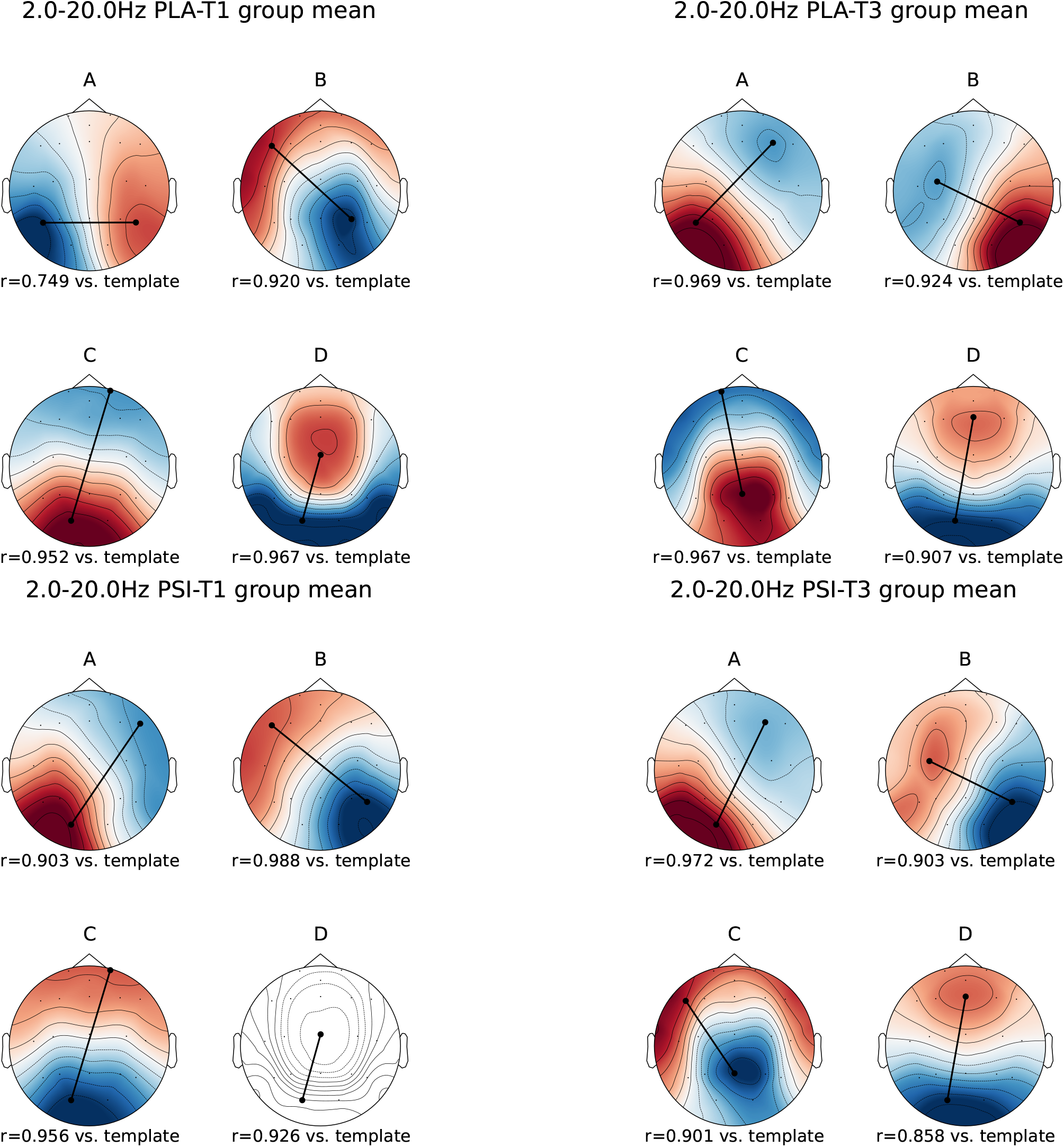
Condition specific topographies of microstates **A, B, C**, and **D** as found by the microstate algorithm for 2–20 Hz bandwidth. Top row: mean maps for placebo T1 (left) and placebo T3 (right); bottom row: mean maps for psilocybin T1 (left) and psilocybin T3 (right). The correlations (*r*) below the plots signify spatial correlation (cf. eq. 2) between each map and template maps provided by Koenig et al. [37].

**Figure 10:**
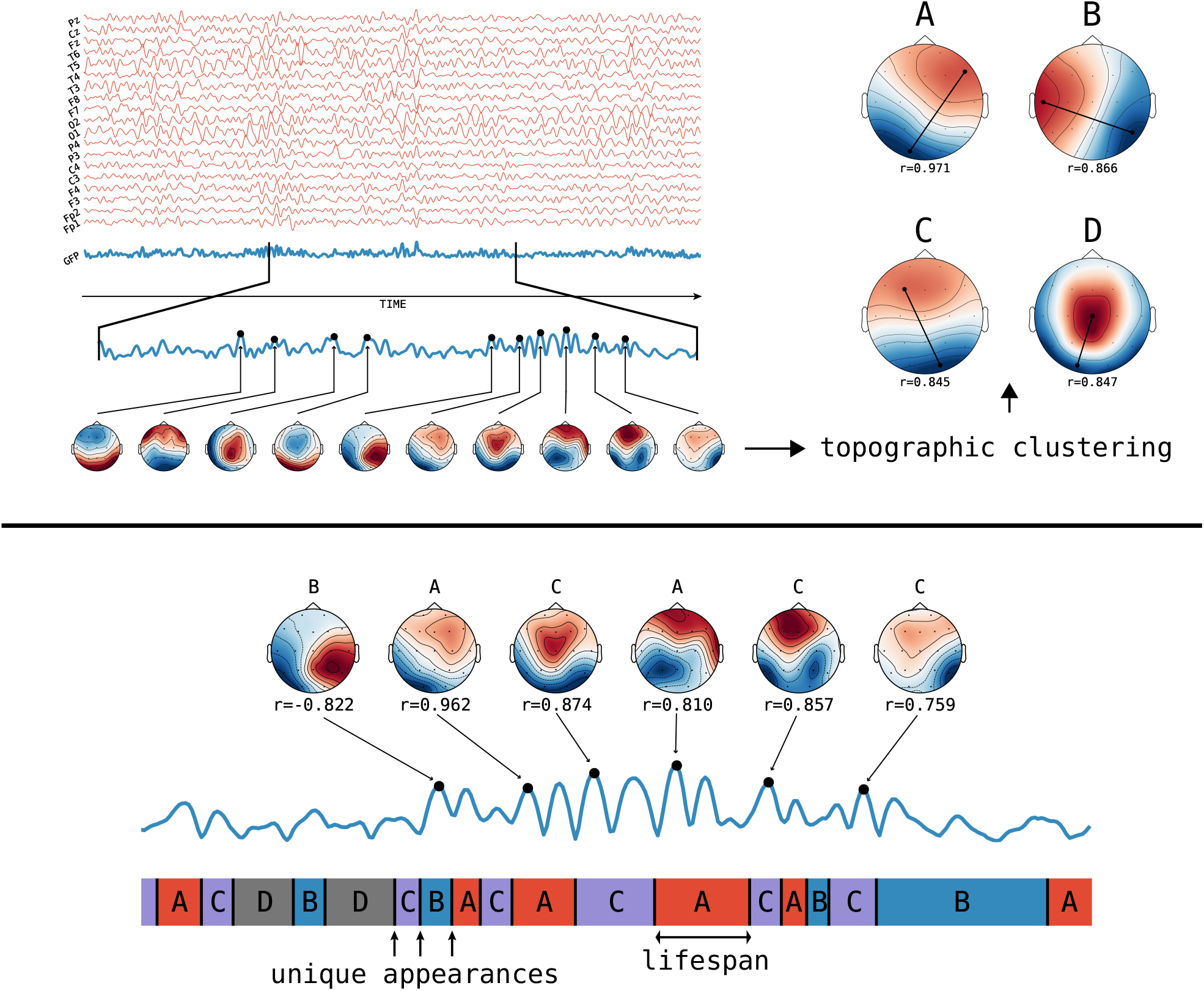
Schematic of the method of microstate analysis and extraction of features of interest from the microstate time series. Adapted from [36]. The GFP (in blue) is calculated at each instant of the multichannel EEG recording (various channels in red). Peaks of the GFP curve represent moments of highest signal-to-noise ratio. At peaks of the GFP curve, the potential recorded at each electrode of the multichannel signal is plotted onto a map of the channel array. This collection of maps is entered into a microstate algorithm (topographic clustering), yielding a small number of canonical maps (**the microstates**) that explain a large proportion of the global topographic variance. Four topographies are repeatedly found using this method; these maps are labeled A, B, C, or D in the figure. Connected dots indicate points of maximum or minimum recorded electric potential. Then, the original maps at peaks of the GFP curve are assigned to a microstate class A, B, C, or D based on the degree of correlation with the microstate maps. This reassignment results in a representation of the original multichannel data as an alternating series of microstates A, B, C, and D. A microstate is considered dominant in the time during which all successive original maps are assigned to the same microstate class. Each period of dominance is considered a unique appearance of a microstate. The frequency of a microstate is the number of unique appearances per second. The coverage of each microstate is the fraction of total recording time that each microstate is dominant. Note: this schematic illustrates the canonical four-microstate solution; in the present study, three microstates (A–C) are identified as optimal in the primary analysis.

**Figure 11:**
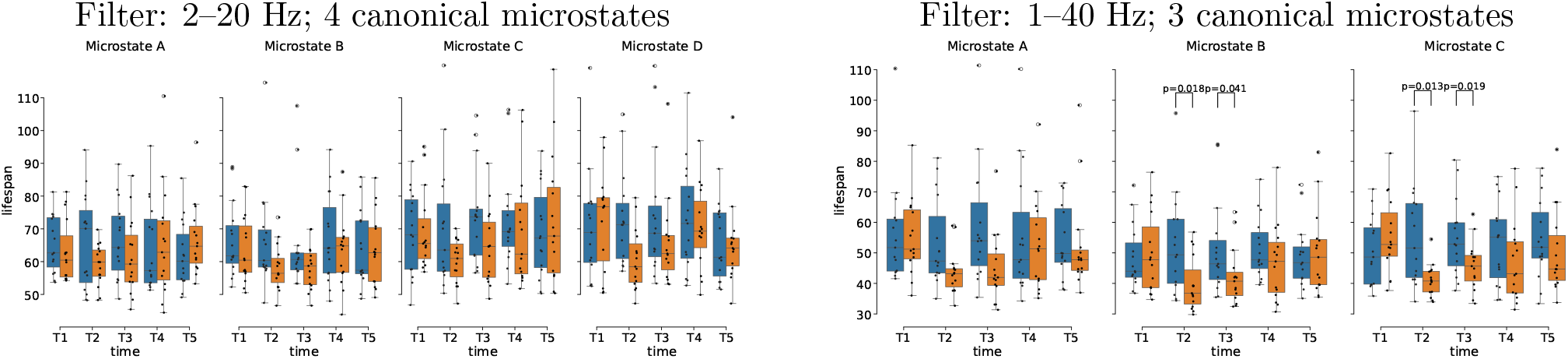
Microstate *×* time interaction for average microstate lifespan in ms. For each filtering paradigm (left: 2–20 Hz; right: 1–40 Hz). Significant differences between placebo and psilocybin condition are marked with vertical lines and their respective *p-values* based on *pairwise t-test* post hoc (preceded by repeated-measures ANOVA), corrected for multiple comparisons using *Benjamini-Hochberg* FDR procedure. For tabulated results see Table 1.

**Figure 12:**
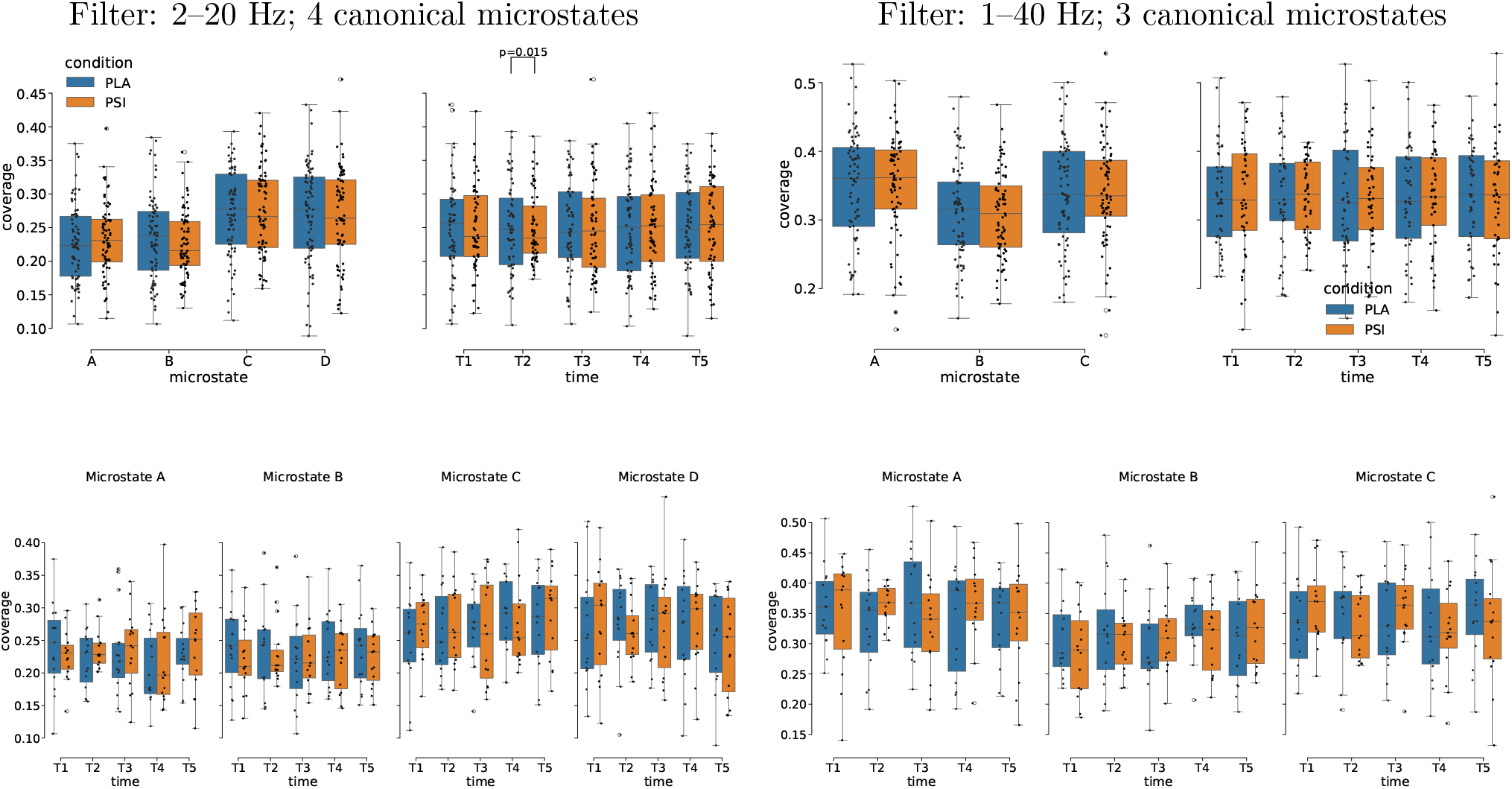
Box plots of total microstate coverage as ratios summing to 1. For each filtering paradigm (left: 2–20 Hz; right: 1–40 Hz), coverage is grouped by microstate (left sub-panel) and time (right sub-panel), with their interaction shown in the bottom row. A significant difference between conditions was found at *T2* in the 2–20 Hz bandwidth (*p* = 0.015, pairwise t-test, Benjamini-Hochberg FDR corrected).

## 4. Discussion

The rationale for analyzing both three- and four-microstate solutions across two frequency bands is detailed in the Methods section. Both approaches yielded qualitatively consistent conclusions across the main outcome measures, supporting the robustness of our findings.

GFP peak density was significantly elevated at T2 and T3 (and at T4 in the 1–40 Hz analysis; Figure 1), corresponding to peak subjective effects. Increased GFP peak density may reflect more frequent transitions between moments of high signal-to-noise ratio and, by extension, more rapid changes in global topographic configurations. This finding aligns with a large body of work showing that psychedelics increase neural signal diversity and entropy, as quantified by Lempel-Ziv complexity and related metrics in EEG and MEG [14, 21, 44, 19, 45]. Computational modeling further supports this view, demonstrating that whole-brain models can reproduce entropy increases under psychedelics through decreased synaptic conductance [46]. Multimodal studies further show that increases in electrophysiological entropy covary with alterations in large-scale functional connectivity [47, 48]. In this context, GFP peak density may provide a complementary, topography-based measure of enhanced dynamical richness under psilocybin.

The most robust microstate-level effect was a pronounced reduction in microstate lifespan at T2 and T3 (Figure 3, Table 1), indicating reduced temporal stability and faster switching between global field configurations. This destabilization is consistent with convergent evidence reviewed in the Introduction, including reductions in alpha and beta power under psychedelics [17, 29, 16, 49] and disruption of large-scale resting-state networks [50, 47]. Notably, microstate statistics are substantially determined by the spectral and autocorrelation structure of the underlying EEG [35, 51], suggesting the observed lifespan reductions are tightly linked to psilocybin-induced spectral acceleration rather than representing a fully independent topographic phenomenon.

The temporal profile of microstate changes closely followed psilocybin pharmacokinetics. Lifespan reductions peaked at T2 (50–60 min) and T3 (90–100 min), matching peak plasma psilocin levels and maximal subjective effects [29]. This pattern of accelerated state transitions is compatible with previous reports linking psychedelic effects to 5-HT_2A_-mediated mechanisms and large-scale network reorganization [47, 52], and with recent evidence for time-dependent alterations in sensory and cognitive processing [53]. Microstate frequency of occurrence increased at T2 and T3 (Figure 4, Table 2), complementing lifespan reductions and consistent with accelerated state transitions. Shorter microstate durations necessarily imply more frequent state onsets within a fixed recording window. Similar acceleration of brain state dynamics has been reported in analyses of metastable topographic sequences and time-resolved EEG under DMT [54, 55, 56], and spontaneous switching between functional connectivity states has been linked to cognitive performance [57]. Together with complexity-based findings, these results support theoretical models proposing that psychedelics increase the rate at which the brain explores its available state space [58, 12].

In contrast to robust effects on temporal parameters, microstate coverage was largely preserved (Supplementary Figure 12), with only a transient significant difference at *T2* in the 2–20 Hz bandwidth (*p* = 0.015). Coverage reflects the proportion of time spent in each microstate class and thus the relative dominance of distinct global configurations. Its broad stability—which should be interpreted cautiously given N=15, as the study may be underpowered to detect smaller coverage changes—suggests that psilocybin is primarily associated with changes in how brain states are sequenced and how long they persist, rather than which states are accessible. This dissociation is notable given that altered microstate coverage has been reported in psychiatric conditions such as schizophrenia and depression [25, 26]. The preservation of coverage alongside altered temporal dynamics may therefore distinguish the psychedelic state from pathological microstate profiles, consistent with the notion that psychedelics transiently increase dynamical flexibility without fundamentally disrupting the repertoire of large-scale brain states [59, 15].

Of potential translational interest were associations between microstate dynamics, acute subjective experience, and persisting effects, most prominently at T2. Using principal component analysis to reduce dimensionality, we observed significant correlations between microstate-derived components and experiential components at T2 (Figures 6, 7). This suggests that inter-individual variation in microstate reorganization tracks variation in subjective phenomenology during the acute psychedelic state. This finding is consistent with prior reports linking electrophysiological measures to experiential intensity, including oscillatory synchronization and signal diversity metrics [60, 21, 61]. Importantly, the same microstate components were also associated with Persisting Effects Questionnaire scores assessed 28 days post-administration, suggesting that the degree of acute microstate destabilization is associated with the magnitude of longer-term psychological change. Greater microstate destabilization at T2 was associated with stronger persisting positive changes in mood, well-being, and personal meaning. However, clinical validation in patient populations is required before any therapeutic inference can be drawn.

These results complement emerging evidence that acute neural signatures are associated with persisting outcomes following psychedelic exposure [62, 63]. Within theoretical frameworks such as the entropic brain and REBUS (Relaxed Beliefs Under Psychedelics) models, transient destabilization of hierarchical constraints and increased dynamical flexibility are proposed to facilitate psychological change by enabling revision of maladaptive priors [58, 13]. Neuroplasticity potentially induced by psychedelics may, speculatively, provide a mechanistic substrate for such lasting effects [64]. While our findings are consistent with this view, they remain correlational and do not establish causality.

Several limitations warrant consideration. The sample size (N=15), while within the range of psychedelic neuroimaging studies, limits sensitivity to smaller effects and fine-grained analyses. Post-hoc power analysis based on observed effect sizes (*α* = 0.05, *N* = 15) indicated adequate power for condition effects in the primary 1–40 Hz analysis: lifespan (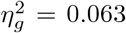, power = 0.68) and frequency of occurrence (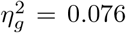, power = 0.78). Power was substantially lower for time effects (power ≤ 0.36) and condition time interactions (power 0.59) in the primary analysis, and across all factors in the secondary 2–20 Hz analysis (maximum power = 0.42), indicating limited sensitivity to smaller or time-resolved effects. An important alternative interpretation of the GFP and lifespan findings is that psilocybin increases heart rate, pupil dilation, and general sympathetic arousal at peak intoxication, which could drive faster EEG fluctuations independently of psychedelic-specific mechanisms. Physiological signals (heart rate, respiration) were not coregistered in the present study, and their potential contribution to microstate dynamics cannot be ruled out; future studies should include concurrent physiological monitoring and regression of cardiovascular artefacts. Crucially, establishing whether microstate dynamics are associated with therapeutic efficacy will require direct testing in clinical populations undergoing psychedelic-assisted treatment. More generally, integration with multimodal imaging, pharmacological manipulations, and computational modeling [47, 52, 48] will be essential for clarifying what specific aspects of neural processing microstates reflect. At present, microstate metrics appear well-suited as pragmatic biomarkers of altered brain dynamics, but their functional interpretation remains incomplete, constraining mechanistic insight into psychedelic therapy.

## 5. Conclusions

Psilocybin was associated with a marked reorganization of resting-state EEG microstate dynamics, characterized by an increased number of GFP peaks, reduced microstate lifespan, and higher frequency of occurrence, while microstate coverage was largely preserved. These findings indicate an acceleration and destabilization of large-scale brain dynamics rather than a change in the repertoire of accessible brain states. Importantly, individual differences in microstate dynamics during the acute psychedelic state were associated with both subjective experience and persisting psychological effects, suggesting that microstate dynamics show promise as neural markers of psychedelic-associated alterations in consciousness with potential therapeutic implications, although their mechanistic interpretation remains limited.

## 6. Supplementary information

## 7. Acknowledgments

The authors thank all participants who volunteered for this study, as well as the technical staff at the National Institute of Mental Health (Klecany, Czech Republic) for their assistance with data collection and EEG recordings.

## References

[1] F. X. Vollenweider, K. H. Preller, Psychedelic drugs: Neurobiology and potential for treatment of psychiatric disorders, Nature Reviews Neuroscience 21 (11) (2020) 611–624. doi:10.1038/s41583-020-0367-2.

[2] R. L. Carhart-Harris, M. Bolstridge, J. Rucker, C. M. J. Day, D. Erritzoe, M. Kaelen, M. Bloomfield, J. A. Rickard, B. Forbes, A. Feilding, D. Taylor, S. Pilling, H. V. Curran, D. J. Nutt, Psilocybin with psychological support for treatment-resistant depression: an open-label feasibility study, The Lancet Psychiatry 3 (7) (2016) 619–627. doi:10.1016/S2215-0366(16)30065-7.

[3] R. R. Griffiths, M. W. Johnson, M. A. Carducci, A. Umbricht, W. A. Richards, B. D. Richards, M. P. Cosimano, M. A. Klinedinst, Psilocybin produces substantial and sustained decreases in depression and anxiety in patients with life-threatening cancer: A randomized double-blind trial, Journal of Psychopharmacology 30 (12) (2016) 1181–1197. doi:10.1177/0269881116675513.

[4] K. Davis, F. S. Barrett, D. G. May, M. P. Cosimano, N. D. Sepeda, M. W. Johnson, P. H. Finan, R. R. Griffiths, Effects of psilocybin-assisted therapy on major depressive disorder: A randomized clinical trial, JAMA Psychiatry 78 (5) (2020) 481–489. doi:10.1001/jamapsychiatry.2020.3285.

[5] J. Rush, M. H. Trivedi, S. R. Wisniewski, A. A. Nierenberg, J. W. Stewart, D. Warden, G. Niederehe, M. E. Thase, P. W. Lavori, B. D. Lebowitz, P. J. McGrath, J. F. Rosenbaum, H. A. Sackeim, D. J. Kupfer, J. Luther, M. Fava, Acute and longer-term outcomes in depressed outpatients requiring one or several treatment steps: a STAR*D report, American Journal of Psychiatry 163 (11) (2006) 1905–1917. doi:10.1176/ajp.2006.163.11.1905.

[6] Cipriani, T. A. Furukawa, G. Salanti, A. Chaimani, L. Z. Atkinson, Y. Ogawa, S. Leucht, H. G. Ruhe, E. H. Turner, J. P. T. Higgins, M. Egger, N. Takeshima, Y. Hayasaka, H. Imai, K. Shinohara, A. Tajika, J. P. A. Ioannidis, J. R. Geddes, Comparative efficacy and acceptability of 21 antidepressant drugs for the acute treatment of adults with major depressive disorder: a systematic review and network meta-analysis, The Lancet 391 (10128) (2018) 1357–1366. doi:10.1016/S0140-6736(17)32802-7.

[7] G. M. Goodwin, S. T. Aaronson, O. Alvarez, P. C. Arden, A. Baker, J. C. Bennett, C. Bird, R. E. Blom, C. Brennan, D. Brusch, L. Burke, K. Campbell-Coker, R. Carhart-Harris, J. Cattell, A. Daniel, C. DeBattista, B. W. Dunlop, K. Eisen, D. Feifel, M. Forbes, H. G. Haumann, D. J. Hellerstein, A. I. Hoppe, M. I. Husain, L. A. Jelen, J. Kamphuis, J. Kawasaki, J. R. Kelly, R. E. Key, R. Kishon, S. Knatz Peck, G. Knight, M. I. G. C. Koolen, M. Lean, R. W. Licht, J. L. Maples-Keller, J. Mars, L. Marwood, M. C. McElhiney, T. L. Miller, A. Mirow, S. Mistry, T. Mletzko-Crowe, L. N. Modlin, R. E. Nielsen, E. M. Nielson, S. R. Offerhaus, V. O’Keane, T. P’alen’ıček, D. Printz, M. C. Rademaker, A. van Reemst, F. Reinholdt, D. Repantis, J. J. Rucker, S. Rudow, S. G. D. Ruffell, A. J. Rush, R. A. Schoevers, M. Seynaeve, S. Shao, J. C. Soares, M. Somers, S. C. Stansfield, D. Sterling, A. Strockis, J. Tsai, L. Visser, M. Wahba, S. Williams, A. H. Young, P. Ywema, S. Zisook, E. Malievskaia, Single-dose psilocybin for a treatment-resistant episode of major depression, New England Journal of Medicine 387 (18) (2022) 1637–1648. doi:10.1056/NEJMoa2206443.

[8] R. L. Carhart-Harris, G. M. Goodwin, The therapeutic potential of psychedelic drugs: Past, present, and future, Neuropsychopharmacology 42 (11) (2017) 2105–2113. doi:10.1038/npp.2017.84.

[9] L. Roseman, D. J. Nutt, R. L. Carhart-Harris, Quality of acute psychedelic experience predicts therapeutic efficacy of psilocybin for treatment-resistant depression, Frontiers in Pharmacology 8 (2018) 974. doi:10.3389/fphar.2017.00974.

[10] R. R. Griffiths, M. W. Johnson, W. A. Richards, B. D. Richards, U. McCann, R. Jesse, Psilocybin occasioned mystical-type experiences: immediate and persisting dose-related effects, Psychopharmacology 218 (4) (2011) 649–665. doi:10.1007/s00213-011-2358-5.

[11] T. Klučkov’a, M. Nikolič, F. Tylš, V. Viktorin, Č. Vejmola, M. Viktorinov’a, A. Bravermanov’a, R. Androvičov’a, V. Andrashko, J. Korč’ak, P. Zach, K. H’ajkov’a, M. Kuchař, M. Bal’ıkov’a, M. Brunovsk’y, J. Hor’aček, T. P’alen’ıček, The phenomenology of psilocybin’s experience mediates subsequent persistent psychological effects independently of sex, previous experience, or setting, Pharmacological Reports 77 (4) (2024) 1024–1039. doi:10.1007/s43440-025-00742-5.

[12] R. L. Carhart-Harris, D. Erritzoe, T. Williams, J. M. Stone, L. J. Reed, A. Colasanti, R. J. Tyacke, R. Leech, A. L. Malizia, K. Murphy, P. Hobden, J. Evans, A. Feilding, R. G. Wise, D. J. Nutt, Neural correlates of the psychedelic state as determined by fmri studies with psilocybin, Proceedings of the National Academy of Sciences 109 (6) (2012) 2138–2143. doi:10.1073/pnas.1119598109.

[13] R. L. Carhart-Harris, R. Leech, P. J. Hellyer, M. Shanahan, A. Feilding, E. Tagliazucchi, D. R. Chialvo, D. Nutt, The entropic brain: a theory of conscious states informed by neuroimaging research with psychedelic drugs, Frontiers in Human Neuroscience 8 (2014) 20. doi:10.3389/fnhum.2014.00020.

[14] M. M. Schartner, R. L. Carhart-Harris, A. B. Barrett, A. K. Seth, S. D. Muthukumaraswamy, Increased spontaneous meg signal diversity for psychoactive doses of ketamine, lsd and psilocybin, Scientific Reports 7 (1) (2017) 46421. doi:10.1038/srep46421.

[15] E. Tagliazucchi, R. L. Carhart-Harris, R. Leech, D. J. Nutt, D. R. Chialvo, Enhanced repertoire of brain dynamical states during the psychedelic experience, Human Brain Mapping 35 (11) (2014) 5442–5456. doi:10.1002/hbm.22562.

[16] S. D. Muthukumaraswamy, R. L. Carhart-Harris, R. J. Moran, M. J. Brookes, T. M. Williams, D. Erritzoe, B. Sessa, A. Papadopoulos, M. Bolstridge, K. D. Singh, A. Feilding, K. J. Friston, D. J. Nutt, Broadband cortical desynchronization underlies the human psychedelic state, The Journal of Neuroscience 33 (38) (2013) 15171–15183. doi:10.1523/JNEUROSCI.2063-13.2013.

[17] M. Kometer, A. Schmidt, L. Jäncke, F. X. Vollenweider, Activation of serotonin 2a receptors underlies the psilocybin-induced effects on α oscillations, n170 visual-evoked potentials, and visual hallucinations, The Journal of Neuroscience 33 (25) (2013) 10544–10551. doi:10.1523/JNEUROSCI.3007-12.2013.

[18] M. Kometer, A. Schmidt, R. Bachmann, E. Studerus, E. Seifritz, F. X. Vollenweider, Psilocybin biases facial recognition, goal-directed behavior, and mood state toward positive relative to negative emotions through different serotonergic subreceptors, Biological Psychiatry 72 (11) (2012) 898–906. doi:10.1016/j.biopsych.2012.04.005.

[19] J. S. Siegel, S. Subramanian, D. Perry, B. P. Kay, E. M. Gordon, T. O. Laumann, T. R. Reneau, N. V. Metcalf, R. V. Chacko, C. Gratton, C. Horan, S. R. Krimmel, J. S. Shimony, J. A. Schweiger, D. F. Wong, D. A. Bender, K. M. Scheidter, F. I. Whiting, J. A. Padawer-Curry, R. T. Shinohara, Y. Chen, J. Moser, E. Yacoub, S. M. Nelson, L. Vizioli, D. A. Fair, E. J. Lenze, R. Carhart-Harris, C. L. Raison, M. E. Raichle, A. Z. Snyder, G. E. Nicol, N. U. F. Dosenbach, Psilocybin desynchronizes the human brain, Nature 632 (8023) (2024) 131–138. doi:10.1038/s41586-024-07624-5.

[20] Č. Vejmola, F. Tylš, V. Pioreck’a, V. Koudelka, L. Kadeř’abek, T. Nov’ak, T. P’alen’ıček, Psilocin, LSD, mescaline, and DOB all induce broadband desynchronization of EEG and disconnection in rats with robust translational validity, Translational Psychiatry 11 (2021) 506. doi:10.1038/s41398-021-01603-4.

[21] C. Timmermann, L. Roseman, M. Schartner, R. Milliere, L. T. J. Williams, D. Erritzoe, S. Muthukumaraswamy, M. Ashton, A. Bendrioua, O. Kaur, S. Turton, M. M. Nour, C. M. Day, R. Leech, D. J. Nutt, R. L. Carhart-Harris, Neural correlates of the dmt experience assessed with multivariate eeg, Scientific Reports 9 (1) (2019) 16324. doi:10.1038/s41598-019-51974-4.

[22] Lehmann, H. Ozaki, I. Pal, EEG alpha map series: brain micro-states by space-oriented adaptive segmentation, Electroencephalography and clinical neurophysiology 67 (3) (1987) 271–288. doi:10.1016/0013-4694(87)90025-3.

[23] M. Michel, T. Koenig, Eeg microstates as a tool for studying the temporal dynamics of whole-brain neuronal networks: A review, NeuroImage 180 (2017) 577–593. doi:10.1016/j.neuroimage.2017.11.062.

[24] J. Britz, D. Van De Ville, C. M. Michel, Bold correlates of eeg topography reveal rapid resting-state network dynamics, NeuroImage 52 (4) (2010) 1162–1170. doi:10.1016/j.neuroimage.2010.02.052.

[25] Lehmann, P. L. Faber, S. Galderisi, W. M. Herrmann, T. Kinoshita, M. Koukkou, A. Mucci, R. D. Pascual-Marqui, N. Saito, J. Wackermann, G. Winterer, T. Koenig, Eeg microstate duration and syntax in acute, medication-naive, first-episode schizophrenia: a multi-center study, Psychiatry Research: Neuroimaging 138 (2) (2005) 141–156. doi:10.1016/j.pscychresns.2004.05.007.

[26] W. K. Strik, T. Dierks, T. Becker, D. Lehmann, Larger topographical variance and decreased duration of brain electric microstates in depression, Journal of Neural Transmission 99 (1-3) (1995) 213–222. doi:10.1007/BF01271480.

[27] T. Koenig, D. Lehmann, M. C. G. Merlo, K. Kochi, D. Hell, M. Koukkou, A deviant EEG brain microstate in acute, neuroleptic-naive schizophrenics at rest, European Archives of Psychiatry and Clinical Neurosciences 249 (4) (1999) 205–211. doi:10.1007/s004060050088.

[28] Y. Yan, M. Gao, Z. Geng, Y. Wu, G. Xiao, L. Wang, X. Pang, C. Yang, S. Zhou, H. Li, P. Hu, X. Wu, K. Wang, Abnormal eeg microstates in alzheimer’s disease: predictors of β-amyloid deposition degree and disease classification, Geroscience 46 (5) (2024) 4779–4792. doi:10.1007/s11357-024-01181-5.

[29] A. Bravermanov’a, M. Viktorinov’a, F. Tylš, T. Nov’ak, R. Androvičov’a, J. Korč’ak, J. Hor’aček, M. Bal’ikov’a, I. Griškova-Bulanova, D. Danielov’a, P. Vlček, P. Mohr, M. Brunovsk’y, V. Koudelka, T. P’alen’ıček, Psilocybin disrupts sensory and higher order cognitive processing but not pre-attentive cognitive processing—study on p300 and mismatch negativity in healthy volunteers, Psychopharmacology 235 (2) (2018) 491–503. doi:10.1007/s00213-017-4807-2.

[30] V. Viktorin, I. Griškova-Bulanova, A. Voicikas, D. Dojčánová, P. Zach, A. Bravermanová, V. Andrashko, J. Horáček, T. Páleníček, Psilocybin-mediated attenuation of gamma band auditory steady-state responses (assr) is driven by the intensity of cognitive and emotional domains of psychedelic experience, Journal of Personalized Medicine 12 (6) (2022) 1004. doi:10.3390/jpm12061004. URL https://doi.org/10.3390/jpm12061004

[31] Dittrich, The standardized psychometric assessment of altered states of consciousness (ascs) in humans, Pharmacopsychiatry 31 (Suppl 2) (1998) 80–84. doi:10.1055/s-2007-979351.

[32] J. E. Overall, L. E. Hollister, P. Pichot, Major psychiatric disorders. a four-dimensional model, Archives of General Psychiatry 16 (2) (1967) 146–151. doi:10.1001/archpsyc.1967.01730200014003.

[33] R. R. Griffiths, W. A. Richards, U. McCann, R. Jesse, Psilocybin can occasion mystical-type experiences having substantial and sustained personal meaning and spiritual significance, Psychopharmacology 187 (3) (2006) 268–283. doi:10.1007/s00213-006-0457-5.

[34] D. Lehmann, W. Skrandies, Reference-free identification of components of checkerboard-evoked multichannel potential fields, Electroencephalography and clinical neurophysiology 48 (6) (1980) 609–621. doi:10.1016/0013-4694(80)90419-8.

[35] N. Jajcay, J. Hlinka, Towards a dynamical understanding of microstate analysis of M/EEG data, NeuroImage 281 (2023) 120371. doi:10.1016/j.neuroimage.2023.120371.

[36] A. Khanna, A. Pascual-Leone, F. Farzan, Reliability of resting-state microstate features in electroencephalography, PloS one 9 (12) (2014). doi:10.1371/journal.pone.0114163.

[37] T. Koenig, L. Prichep, D. Lehmann, P. V. Sosa, E. Braeker, H. Kleinlogel, R. Isenhart, E. R. John, Millisecond by millisecond, year by year: normative eeg microstates and developmental stages, Neuroimage 16 (1) (2002) 41–48. doi:10.1006/nimg.2002.1070.

[38] P. Milz, P. L. Faber, D. Lehmann, T. Koenig, K. Kochi, R. D. Pascual-Marqui, The functional significance of EEG microstates—associations with modalities of thinking, Neuroimage 125 (2016) 643–656. doi:10.1016/j.neuroimage.2015.08.023.

[39] V. F’erat, M. Seeber, C. M. Michel, T. Ros, Beyond broadband: Towards a spectral decomposition of electroencephalography microstates, Human Brain Mapping 43 (10) (2022) 3047–3061. doi:10.1002/hbm.25834.

[40] M. M. Murray, D. Brunet, C. M. Michel, Topographic ERP analyses: A step-by-step tutorial review, Brain Topography 20 (4) (2008) 249–264. doi:10.1007/s10548-008-0054-5.

[41] V. Piorecka, Č. Vejmola, P. Pešková, M. Piorecký, S. Jiříček, V. Koudelka, I. Grišková-Bulanova, T. Páleníček, Microstate in rats’ EEG: a proof of concept study, Translational Psychiatry 15 (2025) 494. doi:10.1038/s41398-025-03702-y.

[42] Y. Benjamini, D. Yekutieli, The control of the false discovery rate in multiple testing under dependency, Annals of statistics (2001) 1165–1188doi:10.1214/aos/1013699998.

[43] R. D. Pascual-Marqui, C. M. Michel, D. Lehmann, Segmentation of brain electrical activity into microstates: model estimation and validation, IEEE Transactions on Biomedical Engineering 42 (7) (1995) 658–665. doi:10.1109/10.391164.

[44] P. A. M. Mediano, F. E. Rosas, C. Timmermann, L. Roseman, D. J. Nutt, A. Feilding, M. Kaelen, M. L. Kringelbach, A. B. Barrett, A. K. Seth, S. D. Muthukumaraswamy, D. Bor, R. L. Carhart-Harris, Effects of external stimulation on psychedelic state neurodynamics, ACS Chemical Neuroscience 15 (3) (2024) 462–471. doi:10.1021/acschemneuro.3c00289.

[45] C. H. Murray, J. Frohlich, C. J. Haggarty, I. Tare, R. Lee, H. de Wit, Neural complexity is increased after low doses of LSD, but not moderate to high doses of oral THC or methamphetamine, Neuropsychopharmacology 49 (2024) 1120–1128. doi:10.1038/s41386-024-01809-2.

[46] R. Herzog, P. A. M. Mediano, F. E. Rosas, P. Lodder, R. Carhart-Harris, Y. Sanz Perl, E. Tagliazucchi, R. Cofre, A whole-brain model of the neural entropy increase elicited by psychedelic drugs, Scientific Reports 13 (1) (2023) 6244. doi:10.1038/s41598-023-32649-7.

[47] C. Timmermann, L. Roseman, S. Haridas, L. Rosas, L. Luan, C. Kettner, D. Martins, R. Leech, A. Feilding, D. J. Nutt, R. L. Carhart-Harris, Human brain effects of DMT assessed via EEG-fMRI, Proceedings of the National Academy of Sciences 120 (13) (2023) e2218949120. doi:10.1073/pnas.2218949120.

[48] S. P. Singleton, C. Timmermann, A. I. Luppi, E. Eckernäs, L. Roseman, R. L. Carhart-Harris, A. Kuceyeski, Network control energy reductions under DMT relate to serotonin receptors, signal diversity, and subjective experience, Communications Biology 8 (2025) 78. doi:10.1038/s42003-025-08078-9.

[49] C. H. Murray, I. Tare, C. Perry, M. Malina, R. J. Lee, H. de Wit, Low doses of LSD reduce broadband oscillatory power and modulate event-related potentials in healthy adults, Psychopharmacology 239 (2021) 1861–1871. doi:10.1007/s00213-021-05991-9.

[50] J. J. Gattuso, D. Perkins, S. Ruffell, A. J. Lawrence, D. Hoyer, L. Jacobson, C. Timmermann, J. Sarris, J. R. McCorvy, S. Pagni, B. J. Hibicke, J. Nichols, D. E. Nutt, Default mode network modulation by psychedelics: A systematic review, International Journal of Neuropsychopharmacology 26 (3) (2022) 155–188. doi:10.1093/ijnp/pyac074.

[51] R. D. Pascual-Marqui, K. Kochi, T. Kinoshita, On the relation between EEG microstates and cross-spectra (Aug. 2022). arXiv:2208.02540.

[52] F. Tylš, Č. Vejmola, V. Koudelka, M. Yamamotov’a, M. Šťastn’y, T. Nov’ak, M. Brunovsk’y, J. Hor’aček, Underlying pharmacological mechanisms of psilocin-induced broadband desynchronization and disconnection of EEG in rats, Frontiers in Neuroscience 17 (2023) 1152578. doi:10.3389/fnins.2023.1152578.

[53] M. Nikoli’c, P. A. M. Mediano, T. Froese, D. Reydellet, T. P’alen’ıček, Psilocybin alters brain activity related to sensory and cognitive processing in a time-dependent manner, medRxivPreprint (2024). doi:10.1101/2024.09.09.24313316.

[54] M. Niedernhuber, D. Suay, M. Mueller, M. Scheidegger, N,N-dimethyltryptamine and harmine formulation shifts metastable topography sequences in the cortex, bioRxivPreprint (2025). doi:10.64898/2025.12.04.692302.

[55] E. Lewis-Healey, C. Pallavicini, F. Cavanna, T. D’Amelio, L. A. De La Fuente, D. Copa, S. Müller, N. Bruno, E. Tagliazucchi, T. Bekinschtein, Time-resolved neural and experience dynamics of medium- and high-dose N,N-dimethyltryptamine, Journal of Cognitive Neuroscience 38 (6) (2025) 1244–1263. doi:10.1162/JOCN.a.2423.

[56] C. Pallavicini, F. Cavanna, F. Zamberlan, L. de la Fuente, L. Ilksoy, C. Perl, C. Arias, M. Romero, R. L. Carhart-Harris, C. Timmermann, E. Tagliazucchi, Neural and subjective effects of inhaled N,N-dimethyltryptamine in natural settings, Journal of Psychopharmacology 35 (4) (2021) 406–420. doi: 10.1177/0269881120981384.

[57] J. Cabral, D. Vidaurre, P. Marques, R. Magalhães, P. Silva Moreira, J. Miguel Soares, G. Deco, N. Sousa, M. L. Kringelbach, Cognitive performance in healthy older adults relates to spontaneous switching between states of functional connectivity during rest, Scientific Reports 7 (2017) 5135. doi:10.1038/s41598-017-05425-7.

[58] R. L. Carhart-Harris, K. J. Friston, REBUS and the anarchic brain: Toward a unified model of the brain action of psychedelics, Pharmacological Reviews 71 (3) (2019) 316–344. doi:10.1124/pr.118.017160.

[59] L.-D. Lord, P. Expert, S. Atasoy, L. Roseman, K. Rapuano, R. Lambiotte, D. J. Nutt, G. Deco, R. L. Carhart-Harris, M. L. Kringelbach, J. Cabral, Dynamical exploration of the repertoire of brain networks at rest is modulated by psilocybin, NeuroImage 199 (2019) 127–142. doi:10.1016/j.neuroimage.2019.05.060.

[60] M. Kometer, T. Pokorny, E. Seifritz, F. X. Vollenweider, Psilocybin-induced spiritual experiences and insightfulness are associated with synchronization of neuronal oscillations, Psychopharmacology 232 (19) (2015) 3663–3676. doi:10.1007/s00213-015-4026-7.

[61] E. Tagliazucchi, F. Zamberlan, F. Cavanna, L. A. de la Fuente, C. Romero, Y. Sanz Perl, C. Pallavicini, Baseline power of theta oscillations predicts mystical-type experiences induced by DMT in a natural setting, Frontiers in Psychiatry 12 (2021) 720066. doi:10.3389/fpsyt.2021.720066.

[62] K. M. Godfrey, B. Weiss, X. Zhang, M. Spriggs, J. Peill, T. Lyons, R. L. Carhart-Harris, D. Erritzoe, An investigation of acute physiological and psychological moderators of psychedelic-induced personality change among healthy volunteers, Neuroscience Applied 4 (2024) 104092. doi:10.1016/j.nsa.2024.104092.

[63] J. Vohryzek, J. Cabral, L.-D. Lord, H. M. Fernandes, L. Roseman, D. J. Nutt, R. L. Carhart-Harris, G. Deco, M. L. Kringelbach, Brain dynamics predictive of response to psilocybin for treatment-resistant depression, Brain Communications 6 (2) (2024) fcae049. doi:10.1093/braincomms/fcae049.

[64] S. F. Grieco, E. Castrén, G. M. Knudsen, A. C. Kwan, D. E. Olson, Y. Zuo, T. C. Holmes, X. Xu, Psychedelics and neural plasticity: Therapeutic implications, Journal of Neuroscience 42 (45) (2022) 8439–8449. doi:10.1523/JNEUROSCI.1121-22.2022.

